# Unravelling the association between non-essential elements and redox homeostasis in wild roe deer

**DOI:** 10.64898/2026.11.17.767290

**Authors:** Amandine Herrada, Benjamin Rey, Jean-François Lemaître, Emmanuelle Gilot-Fromont, Marie-Laure Delignette-Muller, Gilles Bourgoin, François Débias, Paul Revelli, Rebecca Garcia, Sonia Saïd, Maryline Pellerin, Nadia Crini, Clémentine Fritsch, Renaud Scheifler, Jean-Michel Gaillard, Pauline Vuarin

## Abstract

Redox homeostasis, i.e. the balance between oxidative damage and antioxidant defences, is crucial for overall physiological function. Disrupting this balance may lead to oxidative stress. Non-essential elements (e.g. lead, cadmium) exhibit toxicity even at low concentrations and can increase the production of pro-oxidant molecules and impair antioxidant mechanisms. While the adverse effects of acute exposure are well documented, little is known about the impact of chronic, low-level exposure in the wild. Here, we investigated the relationship between non-essential element exposure and redox status in two roe deer populations (*Capreolus capreolus*), while accounting for the potential modulation by essential elements (e.g. selenium, zinc) and local weather conditions. We measured the concentrations of nine non-essential and thirteen essential elements in hair, and assessed the oxidative status through seven biomarkers in blood samples (*n* = 649 samples collected between 2016 and 2019). Our findings revealed an elevation of oxidative damage (malondialdehyde and protein carbonyl levels, and reactive oxygen metabolites), and a depletion of an enzymatic antioxidant (SOD activity) but an increase in total antioxidant capacity, with increasing non-essential element content, especially cadmium and mercury. Essential elements showed limited and biomarker-specific associations with redox status, and modulated some of the reported associations between non-essential elements and oxidative stress biomarkers. Finally, individuals that had recently experienced higher temperatures or rainfall exhibited stronger responses. Overall, our study raises concerns regarding the broader consequences of chronic exposure to non-essential elements for wildlife health.

## 1. Introduction

Redox homeostasis, arising from the precise regulation of the balance between pro-oxidant molecules and antioxidant systems, is a cornerstone of physiological resilience across a broad range of taxa. Redox homeostasis influence the ability of organisms to cope with stressful conditions, supporting functions essential to reproduction and survival (Burton and Jauniaux, 2011; Finkel and Holbrook, 2000; Halliwell and Gutteridge, 2015). Alteration of this balance, typically through over production of pro-oxidants such as reactive oxygen species (ROS), and/or depletion of intrinsic antioxidant systems, result in the detrimental accumulation of oxidative damage to macromolecules (proteins, lipids and nucleic acids) within cellular compartments, a physiological state referred to as oxidative stress (Burton and Jauniaux, 2011; Halliwell and Gutteridge, 2015). Oxidative stress may compromise cellular function and viability, as described in many vertebrate species, and on the long-run might contribute to an acceleration of the ageing process (López-Otín et al., 2023).

Various environmental stressors have been shown to exacerbate ROS production, thereby increasing susceptibility to oxidative stress with possible cascading effects on individual health (e.g. Bilham et al., 2018; Rodríguez-Estival et al., 2016). Among them, organisms are increasingly exposed to various, and often toxic, chemicals stemming from human activities. In particular, non-essential elements, such as arsenic (As), cadmium (Cd), mercury (Hg) or lead (Pb), have, in mammals, no documented biological functions and are toxic even at low levels of exposure (Govind, 2014; Tchounwou et al., 2012). Once released in the environment, these elements can bind to particulate matter and travel tens to hundreds of kilometres from their emission source, before falling out (Azeh Engwa et al., 2019; Shahid et al., 2019). These particles can pass the dermal barrier or be directly inhaled (Azeh Engwa et al., 2019). Additionally, they also deposit in water and soil, from which they are absorbed by organisms, eventually accumulating through food chains (Shahid et al., 2019). On the contrary, essential elements, such as copper (Cu), iron (Fe), selenium (Se) or zinc (Zn), have crucial physiological functions such as metabolism or growth. However, they can exert toxic effects at very high dose of exposure. Actually, both types of elements (essential and non-essential) become harmful when their absorption and storage rates exceed their detoxification or excretion rates (Govind, 2014). They can then interact with cellular components, metabolic enzymes, as well as detoxification and repair pathways (Tchounwou et al., 2012). One key pathway through which these elements exert toxicity is by disrupting redox homeostasis, either directly by increasing ROS production, or indirectly by inhibiting antioxidant systems (Valko et al., 2005). For instance, As, Hg and Pb may bind to crucial antioxidant enzymes such as superoxide dismutase (SOD), impairing their ability to neutralise the harmful pro-oxidant superoxide anions (Kumar et al., 2020).

Most of our knowledge on the adverse health effects of non-essential elements stems from studies on humans and laboratory animals (e.g. in mice and rats, Fan et al., 2020; Xu et al., 2017), even if the field of pollutant exposure has flourished in the past decades (Shore and Rattner, 2001). In the wild, aquatic and avian communities have received the most attention (Rattner, 2009), with studies demonstrating that both acute and chronic exposure to non-essential elements can lead to immunosuppression, endocrine disruption, and behavioural alterations (Boyd, 2010; Govind, 2014; Iavicoli et al., 2009; Saaristo et al., 2018), possibly underpinned by an overall increase in oxidative stress (Isaksson, 2010). However, most existing literature on terrestrial mammals has concentrated on the effects of acute exposure to single elements (e.g. Rodríguez-Estival et al., 2011). Yet, it is more likely that the majority of organisms, including wildlife, are exposed to low levels of multiple mineral elements through their life (e.g. Wirth & Mijal, 2010). Such exposure is also expected to affect individual performance on the long-term, as recently suggested for pesticides (Moreau et al., 2021). In this context, investigating whether low-level exposure to non-essential elements affects wildlife physiology has become particularly relevant.

In addition to the level and duration of exposure, physiological responses to non-essential elements may be modulated by essential element content and other environmental stressors. Essential elements can influence the responses to exposure to non-essential ones. As cofactors of several antioxidant enzymes, these elements may mitigate or exacerbate non-essential element-induced oxidative stress, underscoring their potential modulatory effects (e.g. Valko et al., 2005). Additionally, organisms in the wild are rarely exposed to isolated stressors. For example, variation in weather conditions, and the increasing occurrence of heatwaves, droughts or severe rainfall, have been shown to influence redox homeostasis, as documented in both captive and free-ranging animal populations (see Chainy et al., 2016; Lushchak, 2011; Metcalfe & Alonso-Alvarez, 2010 for reviews). Such stressors may further exacerbate the physiological effects of chemical exposure. Yet, the interactive effects of multiple environmental stressors on oxidative stress remain poorly understood, especially in terrestrial mammals (but see Sokolova & Lannig, 2008; Xiao et al., 2024 for examples on aquatic ectotherms). Accounting for these potential interactions is important for assessing the ecological relevance of contaminant exposure in the wild.

In this study, we focused on the European roe deer (*Capreolus capreolus*), a widespread species known to be highly tolerant to human activities (Bonnot et al., 2013; Linnell and Andersen, 1995). Roe deer are thus potentially exposed to various sources of chemicals including non-essential elements through consumption of contaminated dust, water and/or plants, or through direct inhalation (Shahid et al., 2019). Moreover, as roe deer within a given population occupy small home ranges (Strandgaard, 1972), which widely differ in terms of habitat structure and plant species composition (Pellerin et al., 2010), individuals might differ in terms of both levels and mixture of non-essential element exposure (Demesko et al., 2019). Actually, roe deer are considered as reliable indicators of non-essential element contamination, and more broadly as sentinels for environmental health (Draghi et al., 2023; García et al., 2011).

While a few studies have investigated non-essential element exposure in roe deer (Cappelli et al., 2020; Draghi et al., 2023; Ludolphy et al., 2021), none has examined the implications of such exposure for redox homeostasis nor did they consider its effects on roe deer inhabiting areas distant from major contamination sources, which could still be subject to chronic, low-level exposure. However, previous work highlighted spatial and temporal variations in non-essential element concentrations in roe deer, linking these variations to industries (Cappelli et al., 2020), urbanisation (Draghi et al., 2023) and agriculture (Ludolphy et al., 2021). In other large herbivores, some studies have linked non-essential element levels to redox status. For example, it was reported that increased Pb content was associated with reduced antioxidant levels in red deer (*Cervus elaphus*) living near a mining area in Spain. By contrast, wild boar (*Sus scrofa*) from the same area exhibited the opposite pattern (Rodríguez-Estival et al., 2011), highlighting species-specific responses to elements, and the need for additional field data to fully understand the consequences of non-essential element exposure for redox homeostasis.

Here, we took advantage of the long-term monitoring of two French roe deer populations subject to different ecological contexts. The aim of the study was twofold: i) to decipher the association between non-essential element exposure and redox homeostasis in wild roe deer, and ii) to assess the modulatory effect of essential elements and weather conditions on the influence of non-essential element exposure on oxidative stress. Mineral element concentrations were measured in roe deer hair because it is a non-invasive method and a reliable predictor of element content in the environment (Draghi et al., 2023; Rashed and Soltan, 2005; Tête et al., 2014), though the correlation between hair concentrations and internal concentrations may vary among elements and biological matrices (D’Havé et al., 2006; Tête et al., 2014; Vuarin et al., 2025). We focused on nine non-essential elements whose harmful effects are widely recognised even at low level of exposure. We also measured three biomarkers of oxidative damage, three of antioxidant defences, and one marker of oxidative balance in plasma or in erythrocytes for the latter. We expected individuals with the highest non-essential element burdens to exhibit higher oxidative stress, i.e. an accumulation of oxidative damage on one hand, and a depletion of antioxidant defences on the other hand. Additionally, we hypothesised that these relationships would be modulated by the content in essential elements and weather conditions (i.e. cumulative rainfall and mean ambient temperature the week prior to capture). Specifically, we expected a stronger biomarker response to non-essential element exposure in individuals with minor concentrations of essential elements, as well as a contrasted response in individuals experiencing different weather conditions.

## 2. Materials and Methods

### 2.1. Study populations

The studied roe deer populations are located in two enclosed forests managed by the Office Français de la Biodiversité (OFB): the Territoire d’Etude et d’Expérimentation of Trois-Fontaines (TF – 1360 ha), in north-eastern France (48°43’N, 4°55’E), and the Réserve Biologique Intégrale of Chizé (CH – 2614 ha), in western France (46°05’N, 0°25’W). The TF forest offers a high-quality habitat for roe deer, being characterised by a continental climate and fertile soils (Saïd et al., 2009), although increasing temperatures and drought frequency in spring-summer has decreased habitat quality since 2003 (Gaillard et al., 2013). On the other hand, the CH forest provides a poorer habitat quality for roe deer, and is defined by a temperate oceanic climate, poor soils and frequent summer droughts (Pettorelli et al., 2006). The two populations have been monitored for almost 50 years through a capture-mark-recapture program (since 1975 for TF and 1977 for CH; Gaillard et al., 1993). Captures take place every winter for 10 to 12 days between December and March. Roe deer are captured using drive nets and transported in wood boxes to a handling area. There, sex, age or age-class, hind foot length, and body mass (± 50 g) are determined for all individuals. Roe deer are fitted with ear tags upon their first capture and a sterilised microchip with a unique number is also inserted percutaneously at the back of the neck.

For individuals of known age (i.e. roe deer first captured within their first year of life when age can be assigned with certainty based on the milk tooth sequence; Flerov, 1952), blood and hair samples are also collected. Hair is sampled from the white patch around the rump of the animal and stored in paper envelopes at room temperature. Blood (up to 1 mL/kg with a maximum of 20 mL) is collected from the jugular vein in 20 mL syringes with sterile hypodermic needles (gauge 19, 1.1 x 38 mm), and rapidly transferred to heparinised collection tubes of 4 mL. These tubes are then spun at 2000 g for 10 min and the plasma transferred into sterile cryogenic tubes. Blood cells are washed by adding ∼2 mL of 0.9 % (v/v) NaCl solution, re-spun at 2000 g for 10 min and the buffy coat containing white blood cells is removed. All tubes (containing either plasma or washed erythrocytes) are frozen on site at -80°C in a portable freezer (Telstar SF 8025) until laboratory analyses. At the end of the procedure, roe deer are released at their site of capture. The sample size was determined by the number of individuals captured and for which suitable hair and/or blood samples were available during the 2016-2019 study period. All eligible samples collected as part of the long-term monitoring programme were included in the analyses.

### 2.2. Essential and non-essential element concentrations

The concentrations of 22 essential and non-essential elements were assessed in roe deer hair samples. Roe deer undergo moult twice a year, in May-June and in October (Johnson and Hornby, 1975; Sempéré et al., 1996). Consequently, elements measured in the hair represent what has been accumulated over the hair growth cycle. Since roe deer were captured during winter and the last moult occurred in autumn, we hypothesised that the storage of elements was therefore completed for all individuals by the time of sampling. This prevents any bias regarding the timing of hair collection or potential seasonal effects (Combs et al., 1982).

A total of 649 hair samples, representing 434 individuals captured between 2016 and 2019, were sent to the PEA²t platform of the Chrono-Environnement laboratory (UMR CNRS 6249, Université Marie et Louis Pasteur, France) to measure the concentration of a large set of essential and non-essential elements. Eighteen elements were measured by inductively coupled plasma mass spectrometry (aluminium (Al), As, Cd, cobalt (Co), chromium (Cr), Cu, Fe, Hg, manganese (Mn), molybdenum (Mo), nickel (Ni), Pb, antimony (Sb), strontium (Sr), Se, titanium (Ti), thallium (Tl), Zn), and four elements were measured by inductively coupled plasma atomic emission spectrometry (calcium (Ca), potassium (K), magnesium (Mg), sodium (Na)) following the protocol given in Herrada et al., (2024a). In brief, hair samples were first washed in alternating baths of acetone and ultrapure water using an ultrasonic cleaner to remove any potential external contamination. After drying at 60°C, hair samples were digested in nitric acid and hydrogen peroxide, with reagent volumes adjusted according to sample weight. The resulting solutions were diluted with ultrapure water and filtered. Procedure blanks (acid, hydrogen peroxide and ultra-pure water) and certified reference materials were prepared and analysed using the same methods as the samples. The concentrations of mineral element concentrations in hair are expressed as µg/g dry mass. For values below quantification limits, half of the quantification limit value was assigned for the statistical analyses (USEPA, 2000). The limits of quantification and the proportion of values under these limits are reported for each element in Table S1. As recovery rates were considered satisfactory between 75 % and 125 %, caution should be paid when considering the values for Hg and Na since they exhibited a high (173 %) and a low (46 %) average recovery, respectively (see also Herrada et al., (2024a) for detailed metrology). The following elements were considered as non-essential elements: Al, As, Cd, Hg, Pb, Sb, Sr, Ti, and Tl, since the harmful effects of these elements is widely acknowledged (Tchounwou et al., 2012). The following elements were considered as essential elements: Ca, Co, Cr, Cu, Fe, K, Mg, Mn, Mo, Na, Ni, Se, and Zn according to their established or suspected biological roles (e.g. Zoroddu et al., 2019). Although essential elements can become toxic at high concentrations, this is unlikely to be the case in our studied populations because essential element concentrations are relatively low compared to those recorded in other populations of large herbivores (Herrada et al., 2024a).

### 2.3. Oxidative stress biomarkers

Roe deer redox status was assessed by measuring three complementary biomarkers of oxidative damage in plasma, namely the total amount of reactive oxygen metabolites (d-ROMs test, Diacron International; CV < 6 %), the levels of thiobarbituric Acid Reactive Substances (TBARS), expressed as malondialdehyde (MDA) concentrations and reflecting lipid peroxidation (TBARS Assay kit, Cayman Chemical; CV < 4 %), and the protein carbonyl content as an estimate of oxidative damage to proteins (carbonyl content assay kit, Sigma-Aldrich; CV < 10 %). We also measured two biomarkers of antioxidant defences in plasma, namely the overall non-specific antioxidant capacity of plasma to *in vitro* oxidation (OXY-Adsorbent test, Diacron International; CV < 6 %) and the activity of an enzymatic antioxidant (superoxide dismutase (SOD) activity determination, Sigma-Aldrich; CV < 9 %); as well as two biomarkers of antioxidant defences in erythrocytes, namely a non-enzymatic antioxidant, reduced glutathione (GSH quantification, Sigma-Aldrich), and the reduced-to-oxidised glutathione ratio, GSH:GSSG (oxidised (GSSG) and reduced glutathione quantification, Sigma-Aldrich). All biomarkers were assayed spectrophotometrically (SAFAS MP96 microplate reader, Monaco) following protocols previously described elsewhere (Herrada et al., 2024b), except for the oxidised and reduced glutathione quantification that will be briefly presented below. Raw values of biomarkers are presented in Table S2.

Reduced glutathione (GSH, γ-L-glutamyl-L-cysteinylglycine) is a tripeptide with high antioxidant properties that is converted to its oxidised form (GSSG) in the presence of ROS, making the GSH:GSSG ratio a widely used index of oxidative stress. GSSG is measured after masking GSH, and both GSSG and total glutathione (GSH + GSSG) concentrations are determined spectrophotometrically at 412 nm using DTNB (5,5’-dithiobis (2-nitrobenzoic acid)) and an enzymatic recycling system. GSH concentration is then calculated by subtracting the amount of GSSG from the total amount of glutathione. Levels of GSH are expressed in µmol/L and the GSH:GSSG ratio is in arbitrary units (AU). Measurements were performed in erythrocytes, hence limiting the analysis to individuals for whom sufficient erythrocyte volume was available, and reducing the final sample size. Average intra-plate CV was 5 % for the total glutathione assay and 8 % for the GSSG assay (kit reference number: 38185).

### 2.4. Plasma haemoglobin concentrations

Plasma haemoglobin concentrations were measured to account for the impact of haemolysis on the values of oxidative stress biomarkers (Herrada et al., 2024b). The haemoglobin assay kit (Sigma-Aldrich) utilises the Triton/NaOH method, wherein haems undergo oxidation in alkaline conditions to produce a colorimetric product detectable via spectrophotometry at 400 nm. Levels are determined by referencing against a standard curve and expressed in mg/dL. Average intra-plate CV was less than 10 %.

### 2.5. Weather conditions

Weather data, i.e. daily temperatures and rainfall, were obtained from the Météo-France database (“Météo-France,” 2023), which provides records with a spatial resolution of 8x8 km throughout France since 1958. After targeting each study site (i.e. CH and TF forests), we averaged temperatures and summed rainfall over the week preceding the capture date of each individual, as these values reflect short-term, local weather conditions experienced by each individual prior to sampling. This time frame was chosen to take into account the relatively rapid dynamics of oxidative stress responses (Costantini, 2019). Meanwhile, we are aware that weather conditions may also influence element accumulation in hair. Although not the primary focus of our study, we explored the relationships between non-essential element concentrations in roe deer hair and average weather conditions (i.e. temperature and rainfall) during hair growth (from October to the sampling date) and thus element accumulation. These results, along with a brief discussion, are provided in the supporting information (Fig. S2).

### 2.6. Statistical analyses

All statistical analyses were performed using R (version 4.2.1; R core Team, 2021). Statistical analyses were performed on a maximum of 649 samples corresponding to 434 different individuals. A total of 362 plasma samples were collected in CH (198 females and 164 males) and 287 in TF (147 females and 140 males). Between 2016 and 2019, 87 individuals were captured twice, 46 individuals were captured three times, and 12 individuals were captured every year (i.e. four times). Age ranged from 1 to 14 years.

We first summarised inter-sample variation among all 9 non-essential elements, and among all 13 essential elements, respectively, using Principal Component Analyses (PCA) performed with the *dudi.pca* function from the ade4 R-package (Thioulouse et al., 2018). Because non-essential elements differ from essential ones in that they do not exert physiological or biochemical roles, we performed separate PCAs to derive distinct indices of non-essential and of essential element exposure. This subsequently allowed us to assess whether essential element status or weather conditions modulated the associations between non-essential element exposure and oxidative stress biomarkers.

Element concentrations were first log-transformed to reduce skewness, and then standardised to homogenise variance between variables. We considered all data from 2016 to 2019, including repeated measurements on the same individuals, as well as both populations. Since PCA results remain similar without repeated measurements, and when considering populations separately, respectively (Figs. S2 & S3), this suggests that individuals have no effect on PCA results and that the inclusion of repeated measures does not introduce bias in the analysis. Actually, repeatability analyses indicated low individual-level repeatability of element concentrations across repeated captures (R = 0-0.126), supporting that individual values are not correlated. The advantages of this multivariate approach are three-fold: (i) minimising the number of potential explanatory variables, (ii) discarding the problem of collinearity between explanatory variables by obtaining uncorrelated synthetic variables, and (iii) capturing the covariation between variables across individuals (Thioulouse et al., 2018). To preserve as much variance as possible, we chose to keep the axes with the largest eigenvalues. For both the non-essential and essential element PCAs, we retained the first two principal components. Component selection was based on the scree plots, the proportion of total variance explained by each component, and the interpretability of the resulting axes. For the PCA on non-essential elements, PCT1 and PCT2 explained 46.8% and 17.4% of the variance, respectively, accounting for 64.2% cumulatively. For the PCA on essential elements, PCE1 and PCE2 explained 31.1% and 13.0% of the variance, respectively, accounting for 44.1% cumulatively. Retaining two components in each PCA provided a consistent approach across element groups and allowed to capture a significant proportion of variance in both essential and non-essential element content.

Then, for each oxidative stress biomarker, we fitted a model to analyse the relationships between element concentrations and redox homeostasis. Using linear mixed-effects models performed with the *lmer* function from the lme4 R-package (Bates et al., 2015), each model included as fixed effects PCT1 and PCT2, PCE1 and PCE2, cumulative rainfall (Rain), mean ambient temperature (Temp), as well as their two-way interactions (PCT x PCE, PCT x cumulative rainfall, and PCT x mean ambient temperature). Indeed, we expected the influence of non-essential elements on redox homeostasis to be stronger in individuals with minor concentrations of essential elements, and to differ in individuals that had experienced contrasted local weather conditions prior to capture, such as heavy rainfall and/or harsh winter temperatures. Since factors other than essential element content and recent weather conditions may influence biomarkers values, we also included the additive effects of body mass, age, capture year (2016 to 2019) and plasma haemoglobin levels (except for GSH and GSH:GSSG ratio, which were measured in erythrocytes) as quantitative explanatory variables, and sex and population as qualitative explanatory factors. We effectively expected an increase in oxidative damage and a decrease in antioxidant defences with increasing age, notably considering that alteration of redox homeostasis in response to chemical pollution may be age-dependent (Martin et al., 2024), and with decreasing body mass (Montes et al., 2011). We also expected higher damage and lower defences in males compared to females, and in CH compared to TF due to its lower habitat quality (see Metcalfe & Alonso-Alvarez, 2010 for review). Individual identity (ID) was set as a random effect to consider the non-independence between repeated measurements on the same individuals. Note that entering ID as a random effect resulted in singularity errors for models with MDA, protein carbonyls, SOD, GSH and GSH:GSSG ratio as dependent variables. In that case, a linear model was computed with the *lm* function from the *stats* package, and a single measurement per individual was randomly selected for the analysis. All continuous variables were scaled before analysis. MDA, protein carbonyls, GSH and GSH:GSSG ratio were also log-transformed before scaling to respect normality assumptions. For fawns (< 1 year old), body mass was recalculated as their predicted mass on the 27^th^ of January (i.e. the median date of captures), to account for their growth over winter and thus to ensure comparability between individuals captured at different dates (Douhard et al., 2017). We implemented model selection from the full model based on the Akaike Information Criteria (AICc), using the dredge function from the R-package MuMin (Bartoń, 2013). This method was used to identify the most parsimonious model for each biomarker among the set of candidate models. We first retained all models within 2 ΔAICc (i.e. difference in AICc between a given model and the model with the lowest AICc; Burnham and Anderson, 2002), excluding those with uninformative parameters (Arnold, 2010). We then selected the model with the fewest parameters in accordance with the principle of parsimony. When models had the same number of parameters, we chose the one with the lowest ΔAICc value. The retained models within 2 ΔAICc, as well as the results of the selected models, are presented in Table S3 and S4, respectively.

Normality and homoscedasticity of models’ residuals, as well as linearity and collinearity assumptions, were confirmed by visual inspection with the *check_model* function from the performance R-package (Lüdecke et al., 2021). For each explanatory variable, we present its coefficient (Estimate), as well as its 95 % confidence intervals (95% CI), to assess the direction and magnitude of its relationship with the response variable. The marginal and conditional R² values are also provided to describe the variance explained by the fixed effects and by the retained models, respectively. Finally, because all continuous variables were standardised prior to model fitting, the resulting regression coefficients represent standardised effect sizes. These coefficients express the expected change in the response variable and were used in Figure 4 to allow direct comparisons of the relative magnitude of associations among predictors within the same model.

## 3. Results

### 3.1. Integrative non-essential element exposure indices

The first two principal components of the PCA performed on non-essential elements explained 64.2 % of the total inter-sample variation in non-essential element burdens (Fig. 1). The first axis (PCT1) captured 46.8 % of the variation, and received loadings particularly from Al, As, Pb, Ti, Tl, and to a lesser extent from Sb and Sr. The second axis (PCT2) captured 17.4 % of the inter-sample variation and received loadings mostly from Cd and Hg. Overall, PCA results revealed that individuals with a high PCT1 and/or PCT2 coordinate were characterised by higher non-essential element burden. PCT1 and PCT2 were then interpreted as two independent and integrative indices of non-essential element exposure. Note that individuals from the two populations are not homogeneously distributed along PCA axes, particularly the second one, suggesting higher concentrations of Cd and Hg at TF than at CH.

**Figure 1.**
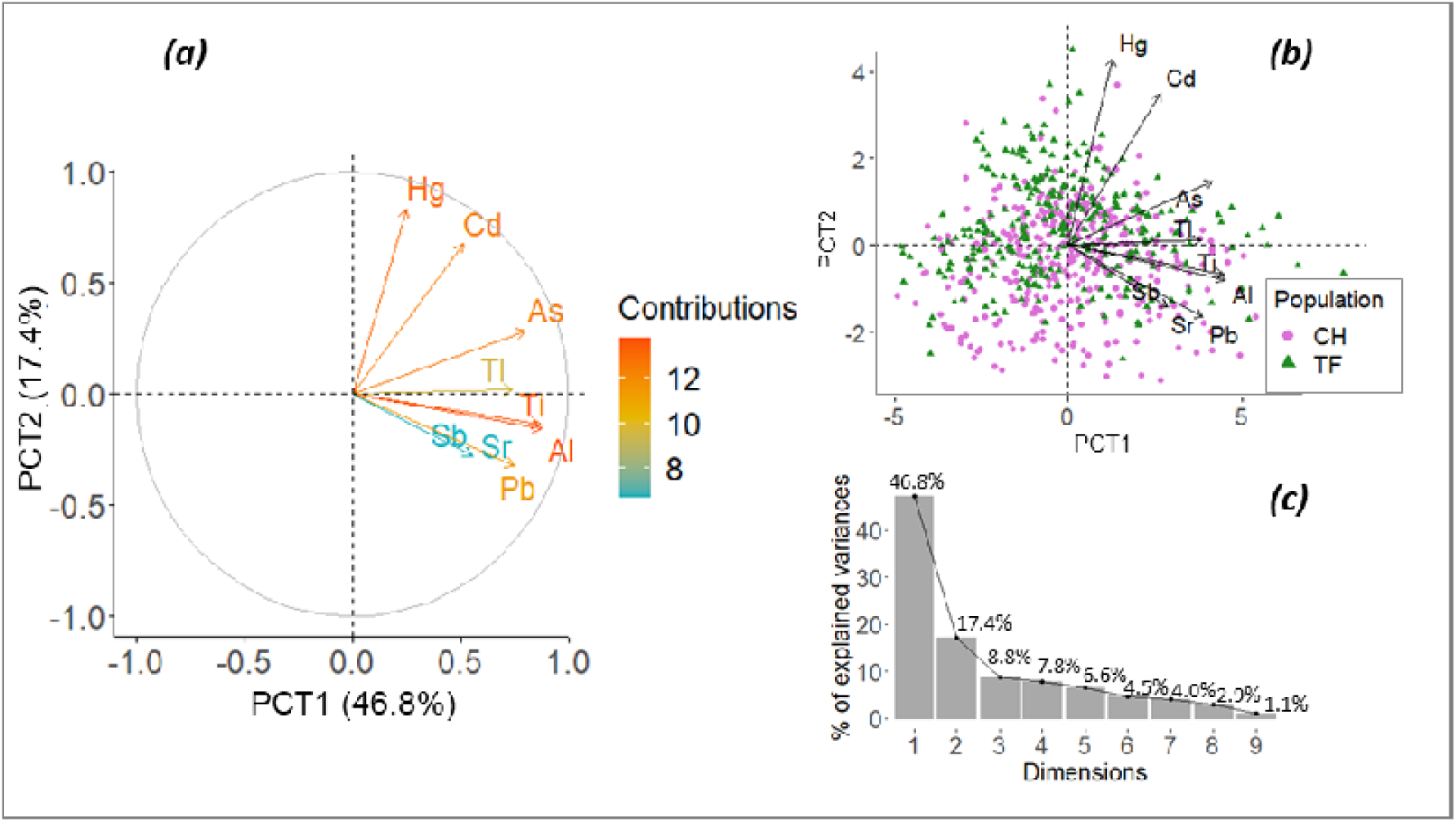
First factorial plan from the Principal Component Analysis (PCA) performed on 9 non-essential elements in roe deer (n = 649). (a) Loadings on the first two axes (PCT1 and PCT2), with colours representing the contribution of the different elements to the first axis, (b) PCA scores of individuals, with green triangles representing individuals from Trois-Fontaines (TF) and lilac dots representing individuals from Chizé (CH), and (c) proportion of inertia contributed by each PC.

### 3.2. Integrative essential element content indices

The first two principal components of the PCA performed on essential elements explained 44.1 % of the total inter-sample variation in essential element concentrations (Fig. 2). The first principal component (PCE1) captured 31.1 % of the variation and received loadings mainly from Ca, Co, Cr, Fe, Mn and Ni. The second principal component (PCE2) captured 13.0 % of the variation and received loadings mostly from K and Na. Overall, the PCA results revealed that individuals with a high PCE1 or PCE2 coordinate were characterised by higher essential element concentrations. These two axes were therefore interpreted as independent and integrative indices of essential element content. Note that individuals from TF also exhibited higher essential element content than those of CH (t-test, t = -4.264, df = 647, *p* < 0.001).

**Figure 2.**
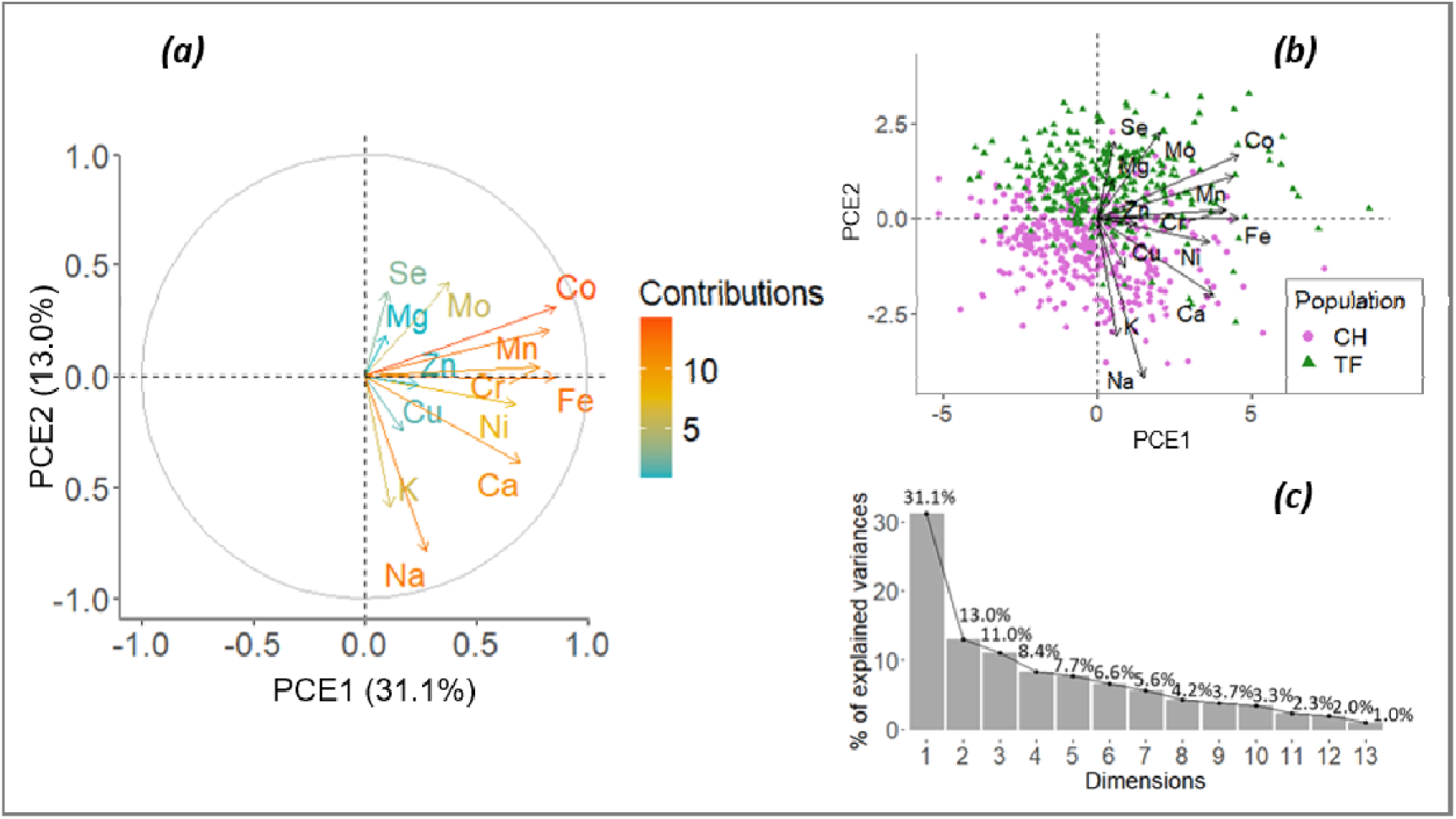
First factorial plan from the Principal Component Analysis performed on 13 essential elements in roe deer (n = 649). (a) Loadings on the first two axes (PCE1 and PCE2), with colours representing the contribution of the different elements to the first axis, (b) PCA scores of individuals, with green triangles representing individuals from Trois-Fontaines (TF) and lilac dots representing individuals from Chizé (CH), and (c) proportion of inertia contributed by each PC.

### 3.3. Relationships between oxidative stress biomarkers, non-essential and essential elements, and weather conditions

#### 3.3.1. Associations between non-essential element exposure and oxidative stress biomarkers

PCT1 coordinates were associated with one antioxidant defence biomarker (OXY), and with the oxidative status marker (i.e. GSH:GSSG ratio). By contrast, PCT2 coordinates were associated with all biomarkers but non-enzymatic antioxidant levels (GSH) (Tables S3,S4; Figs. 3,4). More specifically, higher Cd and Hg burdens were associated with higher oxidative damage on lipids (MDA) and proteins (Carbonyls), as well as higher reactive oxygen metabolite levels (dROMs) (Table S4; Fig. 3a-e). Associations between non-essential elements and antioxidant defences were contrasted as higher element concentrations were associated with higher total antioxidant capacity (OXY), but lower SOD activity (Table S4; Fig. 3f-h). Note that for most markers, effect sizes suggested relatively small associations, as reflected by the standardised coefficient estimates and their 95% confidence intervals (Fig. 4). However, the associations between non-essential elements and biomarkers of oxidative stress may vary according to the essential element content and the weather, as described later (see sections *3.3.2 and 3.3.3*).

**Figure 3.**
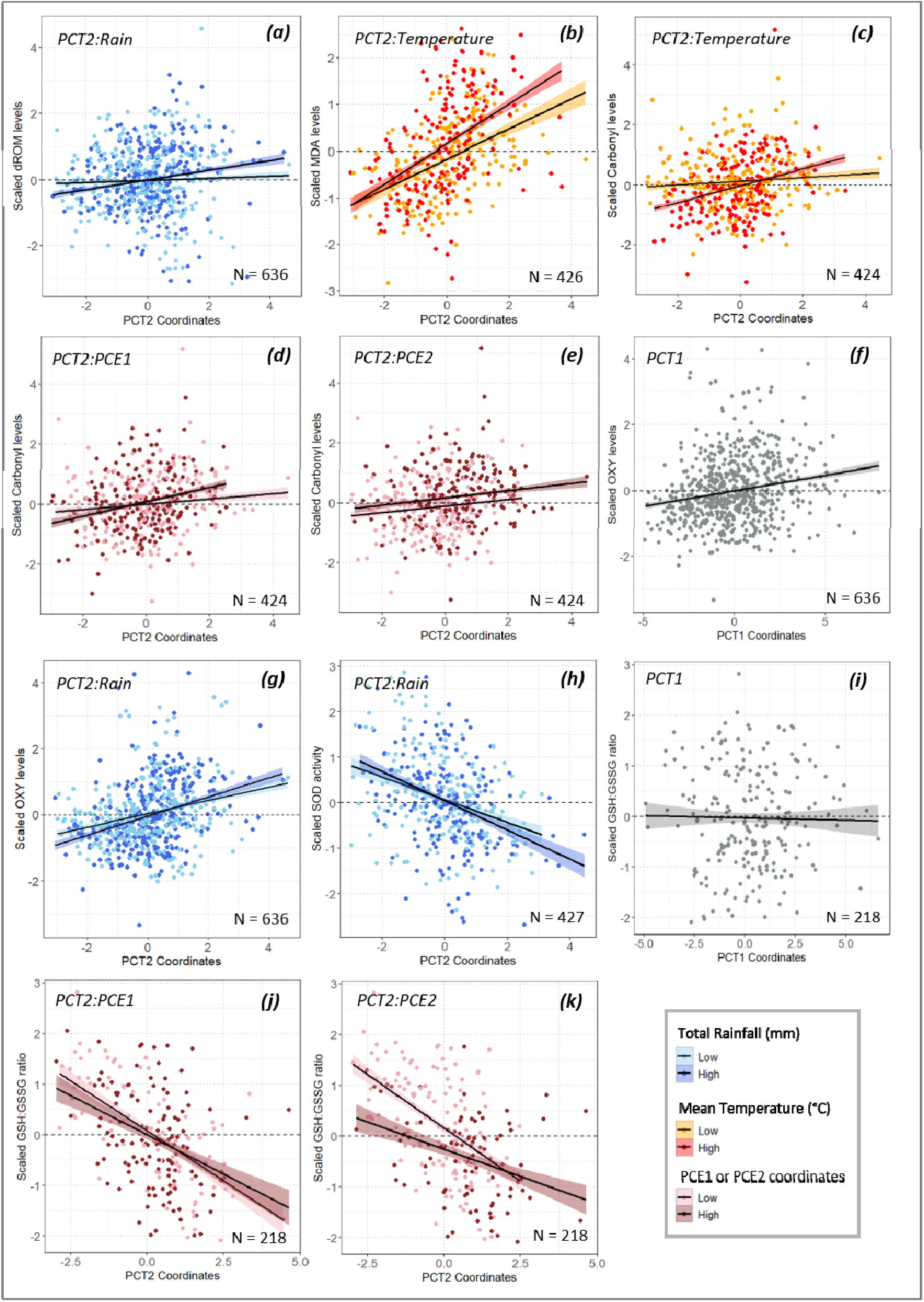
Relationships between individual coordinates on the first or second axis of the PCA performed on non-essential elements (i.e. PCT1 mainly supported by Al, As, Pb, Sb, Sr, Ti and Tl; PCT2 mainly supported by Cd and Hg) and levels of biomarkers of oxidative damage (dROMs, MDA, Carbonyls) and antioxidant defence (SOD, OXY), as wel as oxidative status (GSH:GSSG). Panels show the relationships between PCT1 or PCT2 and (a) dROMs (b) MDA, (c,d,e) Carbonyls, (f,g) OXY, (h) SOD, and (i,j,k) GSH:GSSG ratio, where individuals are coloured according to the mean ambient temperatures or the cumulative rainfall over the week preceding capture, or according to the essentia element content. Points are raw data and lines with shaded areas represent the predictions from linear models with 95 % confidence intervals. For the purpose of graphical representation, individuals were divided into 2 groups based on the weather factor or the essential element indices: low (< median value) versus high (> median value).

**Figure 4.**
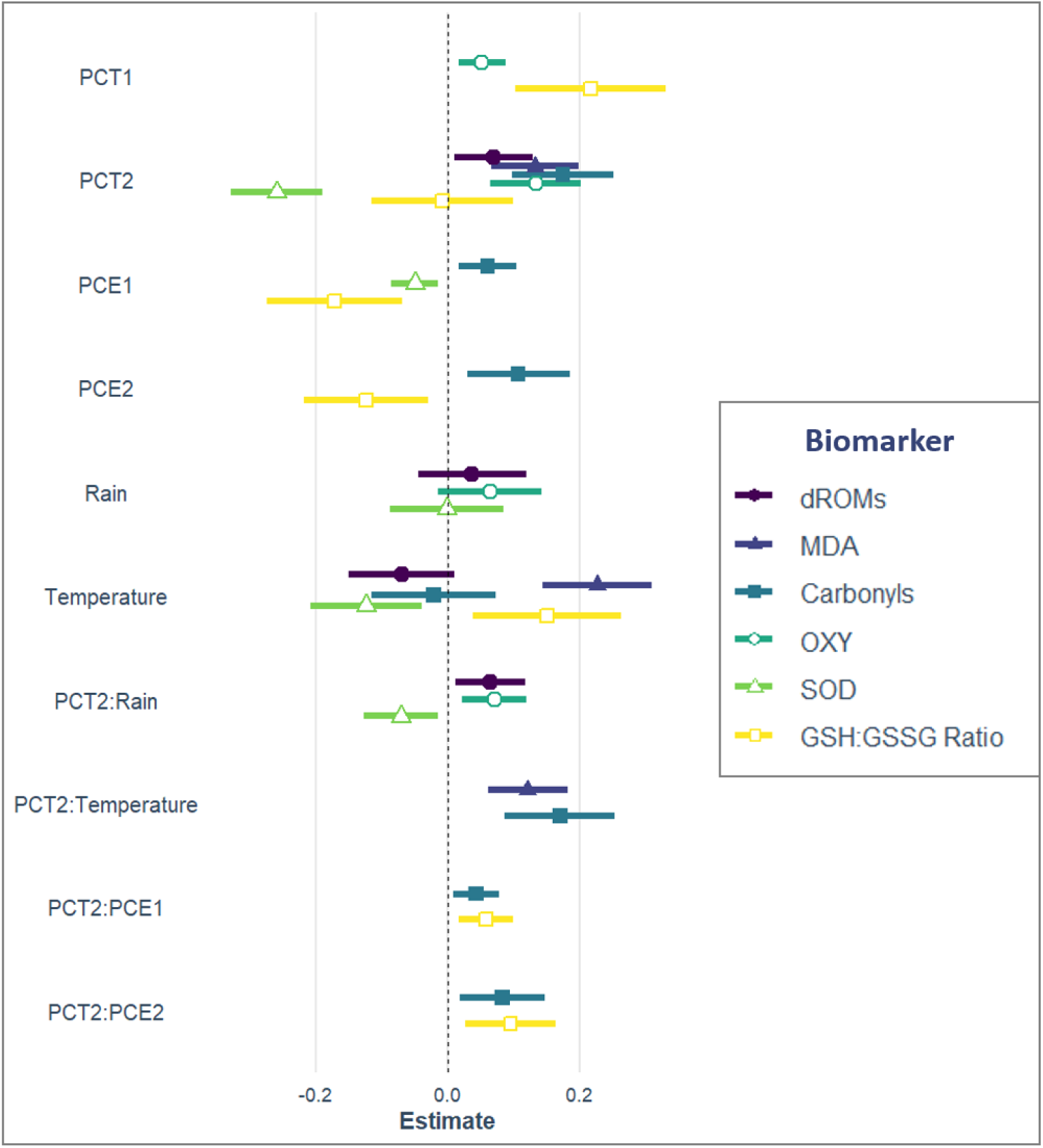
Standardised regression coefficients (β) and 95 % confidence intervals for the association between non-essential element exposure indices (PCT1 and PCT2), essential element content indices (PCE1 and PCE2) and their interaction with non-essential elements (PCT:PCE), cumulative rainfall (Rain) and their interaction with non-essentia elements (PCT:Rain), mean ambient temperature (Temp) and their interaction with non-essential elements (PCT:Temp), and capture date and the biomarkers of oxidative damage (dROMs, MDA, Carbonyls), antioxidant defences (OXY, SOD) and oxidative status (GSH:GSSG). Each oxidative stress biomarker is depicted by a specific colour and symbol.

#### 3.3.2. Associations between essential element content and oxidative stress biomarkers, and their interactions with non-essential ones

Both PCE1 and PCE2 values were positively associated with oxidative damage on proteins (Carbonyls) and negatively associated with the GSH:GSSG ratio. PCE1 was also negatively associated with SOD activity, although the effect size was particularly small (Table S4; Fig. 4). Significant interactions between non-essential and essential elements were found for two biomarkers, indicating that individuals subjected to higher non-essential element burden and higher essential element content showed a greater increase in oxidative damage on proteins and a sharper decrease in the GSH:GSSG ratio, indicating stronger oxidative stress. However, the interaction estimates were small as well (Fig. 4).

#### 3.3.3. Influence of weather conditions and their interactions with non-essential elements

Regarding the effects of weather variables alone, rainfall prior to capture was not clearly associated with any oxidative stress biomarker. In contrast, mean ambient temperatures showed associations with some biomarkers, higher ambient temperatures being associated with higher oxidative damage on lipids (MDA) and a higher GSH:GSSG ratio, but lower SOD activity (Table S4). Significant interactions between non-essential elements and weather conditions were found for five biomarkers, indicating that individuals subjected to higher non-essential element burdens and higher ambient temperatures or higher cumulative rainfall showed a greater increase in oxidative damage (dROMs, MDA, Carbonyls) and antioxidant defences (OXY), as well as lower enzymatic antioxidant activity (SOD) (Table S4; Fig. 3a-c,g,h). This suggests that the relationship between non-essential elements and oxidative stress may be partly modulated by local weather conditions. Effect sizes revealed, however, that these associations were small to almost null and should therefore be interpreted with caution (Fig. 4).

## 4. Discussion

We hypothesised that individuals with substantial non-essential element burdens would be more prone to oxidative stress than those with lower burdens, hence displaying higher oxidative damage and lower antioxidant defences. Our results highlight the existence of relationships between non-essential elements and several oxidative stress biomarkers in roe deer from natural populations. However, these relationships were generally modest in magnitude. Essential elements and weather conditions also appeared to modulate some of the relationships between non-essential elements and oxidative stress biomarkers, although these effects were generally small as well.

### 4.1. The relative contribution of non-essential elements to oxidative stress

Our results provide evidence for associations between exposure to non-essential elements and various biomarkers of oxidative stress in wild roe deer from our studied populations, as shown by the positive relationships between non-essential element concentrations in hair and the accumulation of oxidised lipids and proteins, and to a lesser extent, other derivatives of reactive oxygen metabolites in plasma. The shift in oxidative balance was also evident from the negative correlation between non-essential element concentrations and the SOD activity, a key antioxidant enzyme acting as a major endogenous ROS scavenger. Only the overall non-specific antioxidant capacity of plasma deviated from our prediction, showing an unexpected positive association with non-essential element concentrations. These associations were particularly pronounced with PCT2, an integrative index of exposure to Cd and Hg, which emerged as a predictor for all but two biomarkers. When focusing on PCT1, reflecting the exposure to Al, As, Pb, Ti, Tl, Sb and Sr, significant associations were only detected with the plasma total antioxidant capacity and the antioxidant status. This pattern may reflect efficient antioxidant defences able to cope with the stress induced by exposure to non-essential elements (Isaksson, 2020). The distinct associations between oxidative stress biomarkers (either antioxidant responses or oxidative damage) and PCT1 and PCT2 may reflect the specific physiological consequences of exposure to different non-essential elements. Overall, our findings are particularly compelling as it was recently reported that the concentrations of non-essential elements in hair of roe deer from the two studied populations are relatively low compared to other populations of roe deer and other large ungulates (Herrada et al., 2024a). Our findings therefore suggest that redox homeostasis may respond to variations in exposure to non-essential elements, even at relatively low, environmentally relevant concentrations.

Field studies on the impact of non-essential elements on mammal physiology remain limited. A few studies nevertheless focused on long-term contamination, showing results consistent with ours. For example, Pb exposure has been associated with increased lipid peroxidation in the liver of red deer from a contaminated area (*Cervus elaphus*; Rodríguez-Estival et al., 2011) and decreased SOD activity in the spleen of wild boars from the same area (*Sus scrofa*; Rodríguez-Estival et al., 2013). Total antioxidant capacity (TAC) also increased with increasing Cd and Pb renal concentrations in brown bears (*Ursus arctos*; Lazarus et al., 2024). However, other trends were also evidenced, such as an increase in glutathione content and a decrease in reactive oxygen species with increasing Cd and Pb concentrations in brown bears (Lazarus et al., 2024), as well as a decrease in GSH levels in the spleen of red deer and wild boars with increasing Pb concentrations (Rodríguez-Estival et al., 2013). Note, however, that the above-cited studies measured mineral elements in organs rather than hair. While hair concentrations are a reliable predictor of element content in the environment, and have been used to estimate organ concentrations, correlations between hair and other matrices remain often limited (e.g. D’Havé et al., 2006; Tête et al., 2014) notably in roe deer (Vuarin et al., 2025). Additionally, non-essential element concentrations in roe deer hair reflect accumulation over a few months (e.g. Sempéré et al., 1996), when hair grew and their roots were still connected to the bloodstream (McLean et al., 2009). By contrast, oxidative stress biomarkers measured in blood are transient and reflect circulating levels within a much narrower period. Furthermore, seasonal variation in circulating element concentrations may further contribute to this temporal mismatch and may therefore not be fully captured by hair element concentrations (Pokorny and Ribaric-Lasnik, 2002). Although this temporal mismatch may affect the interpretation of the observed associations, the fact that we found significant relationships between non-essential elements and oxidative stress biomarkers, and that those relationships are mostly of similar direction for markers that represent comparable processes (e.g. oxidative damage), supports that the observed oxidative profiles likely reflect the cumulative effects of chronic exposure.

The fact that almost all plasma and cellular oxidative stress biomarkers were related to hair non-essential element content highlights the broad impact of these elements and indicates that various components of redox homeostasis could be affected. Considering that each biomarker may reflect distinct biochemical processes (Christensen et al., 2015), this also underscores the complexity of the interactions between oxidative stress and mineral elements, and the varying sensitivities of each biomarker to different elements. For example, the bimetallic CuZn-SODs that can be found in the cytosol (SOD1 isozyme) and in the extracellular space (SOD3 isozyme) are directly affected by redox-inactive elements such as As, Cd, Hg, and Pb. Indeed, these elements may interact with the SODs’ cofactors, preventing their normal functioning at least in mammals, which normally consists in the dismutation of superoxide anions into less harmful hydrogen peroxides (H_2_O_2_) (Kumar et al., 2020; Nzengue et al., 2011). This mechanism may therefore play a role in the reduced SOD activity observed in roe deer with relatively elevated non-essential element content. In such case, other endogenous, but also exogenous, antioxidants may help compensate for specific antioxidant decline, resulting in differences between antioxidants’ response to elements (e.g. Losdat et al., 2011). Such mechanisms may explain the observed increase in the overall non-specific antioxidant capacity (OXY) observed in roe deer.

### 4.2. The influence of essential elements in non-essential element-induced oxidative stress

Several essential elements are directly or indirectly involved in maintaining redox homeostasis. They can even partly modulate the effects of non-essential elements on the oxidative balance. For instance, Se is a component of glutathione peroxidase (GPx) and has thus an indirect protective role against oxidative damage, especially from hydrogen peroxide (Soetan et al., 2010; Tapiero et al., 2003). As already mentioned, Cu and Zn are crucial for redox homeostasis since they act as cofactors in SOD1 and SOD3 enzymes, as well as Mn which acts as a cofactor in the mitochondrial SOD2 enzyme (Kumar et al., 2020). Therefore, individuals with substantial concentrations of such essential elements should benefit from their protective effect and be expected to exhibit less pronounced relationships between oxidative stress biomarkers and non-essential elements. However, our analyses did not provide consistent evidence for such a protective effect. In addition, PCE1, mainly driven by Ca, Co, Cr, Fe, Mn and Ni, included elements that exhibit complex relationships with the redox balance, such as Fe and Co that can participate in redox reactions and may become pro-oxidant under certain conditions (Valko et al., 2005). PCE1 and PCE2, which are mainly driven by K and Na, were positively associated with protein carbonyl content and negatively associated with the GSH:GSSG ratio. These associations suggest that variation in essential element profiles may also be related to changes in oxidative stress biomarkers. However, they do not necessarily indicate adverse effects, particularly given the physiological functions of essential elements and the relatively low concentrations measured in our study populations compared with those reported in other large herbivores (Herrada et al., 2024a). Moreover, PCE1 and PCE2 are composite indices of element concentrations in hair rather than direct measures of physiologically available essential elements. Hair concentrations may reflect circulating levels during hair growth, but they do not necessarily reflect the functional pools of essential elements involved in antioxidant defences.

Overall, our results might indicate that the physiological disturbance induced by non-essential elements did not exceed a threshold where compensation mechanisms involving essential elements would be mobilised. We highlighted relationships between most biomarkers and non-essential elements, and it was particularly evident for Cd and Hg. However, while observing these associations, we need to consider that they may not necessarily translate into long-term physiological consequences. It is indeed plausible that the antioxidant repair mechanisms may efficiently counteract oxidative damage. This raises the question of the biological relevance of the observed relations between biomarkers and non-essential elements and would require identifying threshold levels beyond which such exposure would have long-term consequences in terms of ageing and health. Unfortunately, such thresholds are poorly documented and difficult to determine (Lushchak, 2014). Although our analyses do not allow us to identify thresholds beyond which non-essential element exposure would impair redox homeostasis, reporting standardised effect sizes may facilitate comparisons among studies. If future studies consistently report associations in the same direction, and if the magnitude of these associations increases with exposure level, this could help characterise exposure-response relationships and identify the exposure ranges at which the disruption of redox homeostasis would become biologically meaningful.

### 4.3. The influence of weather conditions on element toxicity

Oxidative stress may occur and persist for hours, days or weeks following exposure to a stressor, depending on its nature and intensity. Among potential stressors, weather conditions have received some attention, including extreme temperatures and rainfall (Chainy et al., 2016; Lushchak, 2011; Metcalfe and Alonso-Alvarez, 2010). In our study, warmer local temperatures the week preceding capture were associated with higher oxidative damage to lipids and lower antioxidant capacity, though most of these effects depended on interactions with non-essential element concentrations. No significant effect of cumulative rainfall was detected. Importantly, we observed differences between populations, with individuals from TF exhibiting higher oxidative damage on lipids and lower antioxidant defences than individuals from CH. While this may reflect contrasted environmental conditions between the two study sites, it could also result from differences in element exposure, as we showed higher non-essential element content in TF on average, especially for Cd and Hg.

When examining the influence of weather conditions on non-essential element-induced oxidative stress, our results suggest that it is indeed, albeit modestly, modulated by temperature and rainfall. Specifically, we found that the relationships between non-essential elements and oxidative stress biomarkers were overall amplified under higher local temperatures and higher cumulative rainfall. Although effect sizes were small, these interactions seem consistent with the limited literature on the subject. For example, studies have shown that increased temperatures tend to exacerbate the toxic effects of elements on organisms, notably by intensifying oxidative stress (see Sokolova & Lannig, 2008; Xiao et al., 2024 for reviews on aquatic organisms). Hence, these findings underscore that short-term weather conditions may subtly influence oxidative stress responses to non-essential elements, an effect that should not be overlooked in future field studies.

### 4.4. Limits of the study

Since each oxidative stress biomarker captures a different facet of the redox status, interpretation of results may remain incomplete if biomarkers are analysed separately. For instance, the TBARS assay, which is based on absorbance spectrophotometry and supposedly estimates MDA as a proxy for lipid peroxidation, has well-documented drawbacks. Indeed, since TBA may react non-specifically with substances other than MDA in the sample, it can artifactually generate lipid peroxidation and result in an overestimation of MDA levels (Halliwell & Gutteridge, 2015; Murphy et al., 2022; but see Aguilar Diaz De Leon & Borges, 2020 for an updated use of this method). Other absorbance spectrophotometry-based biomarker assays used in this study might also be criticised for their low reproducibility and non-specificity (e.g. OXY, dROMs; Murphy et al., 2022). Nevertheless, the use of seven oxidative stress biomarkers that reflect different but complementary processes (i.e. lipid and protein oxidation, enzymatic and non-enzymatic defences, overall oxidative status through the GSH:GSSG ratio), should, at least partly, mitigate the above-mentioned issues, and provide an integrative view of the redox status when combined and interpreted together. Most importantly, the results show that the biomarkers follow the expected patterns, and co-vary on a given side of the balance (e.g. all markers of oxidative damage increase with PCT2 values), hence reinforcing the conclusions drawn from the findings of this study.

Since redox reactions can be celland tissue-specific (e.g. Costantini, 2019), we are aware that the measurements of biomarkers in blood (either plasma or erythrocytes) may not always reflect oxidative dynamics elsewhere in the body. Ideally, oxidative stress biomarkers should be measured in other tissues as well (e.g. liver, muscle, kidney, etc.), and ultimately, so should non-essential elements (see section 4.1.). However, this study uses data from a long-term capture-recapture program, where only non-destructive techniques can be applied to collect biological samples. While hair and blood may remain imperfect proxies for internal tissue exposure and oxidative status assessments, they are classically used in ecophysiology studies, making the comparison between studies easy. We also sought to control known biases associated with these matrices, for instance by accounting for the effect of haemolysis on the measurement of plasma biomarkers (Herrada et al., 2024b).

Finally, although several detected effects were statistically significant, their magnitude was often modest, raising the question of their biological relevance. Future studies could evaluate whether such physiological alterations have measurable impacts on individual fitness and demographic trajectories (e.g. Ouyang et al., 2016). It is also important to keep in mind that the reported associations between non-essential element exposure and oxidative stress biomarkers do not imply causal relationships. We cannot rule out that the observed oxidative profiles could be shaped, at least partially, by unmeasured environmental factors (e.g. availability and quality of resources) or individual state (e.g. parasite burden).

## 5. Conclusion

Key results of the present study revealed a clear association of non-essential elements with redox homeostasis in wild roe deer, despite relatively low-level exposure. There was, however, only a moderate support for a modulation of these effects by essential elements and by local weather conditions. Our study therefore adds valuable data to the scattered knowledge on the relationship between exposure to non-essential elements and oxidative stress in natural populations. It also improves our understanding of the potential, although overlooked, impact of low-level element exposure on wildlife physiology. Finally, as oxidative stress has been reported to promote ageing, to impact health and ultimately survival (Finkel and Holbrook, 2000; Halliwell and Gutteridge, 2015), the next step would be to analyse whether non-essential element-induced disruption of redox homeostasis actually contributes to these processes. Only a handful of studies have suggested an implication of toxic chemical exposure for population dynamics in the wild, focusing on marine birds (e.g. Goutte et al., 2014, 2018). Our study thus calls for greater attention to the physiological costs of chronic non-essential element exposure in terrestrial animals.

## Acknowledgements

We are grateful to all the technicians and volunteers who participated in the captures and data collection in Chizé and Trois-Fontaines forests. We also would like to thank Louise Cheynel, Lou Decosne, Alexia Gache and Yoan Regnaud for their participation in the assessment of oxidative stress biomarkers.

## CRediT authorship contribution statement

Amandine Herrada: Conceptualization, Formal analysis, Writing – original draft, Writing – review & editing. Benjamin Rey: Conceptualization, Data curation, Supervision, Writing – review & editing. Jean-François Lemaître: Conceptualization, Writing – review & editing. Emmanuelle Gilot-Fromont: Writing – review & editing. Marie-Laure Delignette-Muller : Formal analysis, Writing – review & editing. Gilles Bourgoin : Funding acquisition, Writing – review & editing. François Débias: Methodology. Paul Revelli: Methodology. Rebecca Garcia: Methodology. Sonia Saïd: Funding acquisition, Writing – review & editing. Maryline Pellerin: Project administration. Nadia Crini: Methodology, Writing – review & editing. Clémentine Fritsch: Methodology, Writing – review & editing. Renaud Scheifler: Methodology, Writing – review & editing. Jean-Michel Gaillard: Supervision, Project administration, Writing – review & editing. Pauline Vuarin: Conceptualization, Data curation, Funding acquisition, Supervision, Writing – review & editing.

## Ethical notes

Game captures were conducted in accordance with European and French laws. The experiment was designed to minimise animal stress and handling time, and to ensure animal welfare, as defined in the guidelines for the ethical use of animals in research. Captures are carried out in accordance with the French Environmental Code (Art. R421-15 to 421-31 and R422-92 to 422-94-1) and approved by the Prefecture of Paris (agreement n°2009-14, n°2013-118, n°2019-02-19-003). The Trois-Fontaines population is part of the long-term Studies in Ecology and Evolution (SEE-Life) program of the CNRS.

## Data accessibility

The datasets and codes supporting the findings of this study are available via a Zenodo repository: https://doi.org/10.5281/zenodo.24314844.

## References

Aguilar Diaz De Leon, J., Borges, C.R., 2020. Evaluation of oxidative stress in biological samples using the thiobarbituric acid reactive substances assay. J Vis Exp. 10.3791/61122

Arnold, T.W., 2010. Uninformative parameters and model selection using Akaike’s Information Criterion. The Journal of Wildlife Management 74, 1175–1178. 10.1111/j.1937-2817.2010.tb01236.x

Azeh Engwa, G., Udoka Ferdinand, P., Nweke Nwalo, F., N. Unachukwu, M., 2019. Mechanism and health effects of heavy metal toxicity in humans, in: Karcioglu, O., Arslan, B. (Eds.), Poisoning in the Modern World - New Tricks for an Old Dog? IntechOpen. 10.5772/intechopen.82511

Bartoń, K., 2013. MuMIn: Multi-model inference. R package version 1.10.0.

Bates, D., Mächler, M., Bolker, B., Walker, S., 2015. Fitting linear mixed-effects models using **lme4**. Journal of Statistical Software 67. 10.18637/jss.v067.i01

Bilham, K., Newman, C., Buesching, C.D., Noonan, M.J., Boyd, A., Smith, A.L., Macdonald, D.W., 2018. Effects of weather conditions on oxidative stress, oxidative damage, and antioxidant capacity in a wild-living mammal, the European badger (*Meles meles*). Physiological and Biochemical Zoology 91, 987–1004. 10.1086/698609

Bonnot, N., Morellet, N., Verheyden, H., Cargnelutti, B., Lourtet, B., Klein, F., Hewison, A.J.M., 2013. Habitat use under predation risk: Hunting, roads and human dwellings influence the spatial behaviour of roe deer. European Journal of Wildlife Research 59, 185–193. 10.1007/s10344-012-0665-8

Boyd, R.S., 2010. Heavy metal pollutants and chemical ecology: Exploring new frontiers. Journal of Chemical Ecology 36, 46–58. 10.1007/s10886-009-9730-5

Burnham, K.P., Anderson, D.R., 2002. Model selection and multimodel inference: A practical information-theoretic approach, 2nd ed. ed. Springer, New York.

Burton, G.J., Jauniaux, E., 2011. Oxidative stress. Best Practice & Research Clinical Obstetrics & Gynaecology, Placental Bed & Maternal - Fetal Disorders 25, 287–299. 10.1016/j.bpobgyn.2010.10.016

Cappelli, J., Frasca, I., García, A., Landete-Castillejos, T., Luccarini, S., Gallego, L., Morimando, F., Varuzza, P., Zaccaroni, M., 2020. Roe deer as a bioindicator: Preliminary data on the impact of the geothermal power plants on the mineral profile in internal and bone tissues in Tuscany (Italy). Environmental Science and Pollution Research 27, 36121–36131. 10.1007/s11356-020-09708-x

Chainy, G.B.N., Paital, B., Dandapat, J., 2016. An overview of seasonal changes in oxidative stress and antioxidant defence parameters in some invertebrate and vertebrate species. Scientifica 2016, 6126570. 10.1155/2016/6126570

Christensen, L.L., Selman, C., Blount, J.D., Pilkington, J.G., Watt, K.A., Pemberton, J.M., Reid, J.M., Nussey, D.H., 2015. Plasma markers of oxidative stress are uncorrelated in a wild mammal. Ecology and Evolution 5, 5096–5108. 10.1002/ece3.1771

Combs, D., Goodrich, R., Meiske, J., 1982. Mineral concentrations in hair as indicators of mineral status: A review. Journal of animal science 54, 391–8. 10.2527/jas1982.542391x

Costantini, D., 2019. Understanding diversity in oxidative status and oxidative stress: The opportunities and challenges ahead. Journal of Experimental Biology 222, jeb194688. 10.1242/jeb.194688

Demesko, J., Markowski, J., Demesko, E., Słaba, M., Hejduk, J., Minias, P., 2019. Ecotype variation in trace element content of hard tissues in the European roe deer (*Capreolus capreolus*). Arch Environ Contam Toxicol 76, 76–86. 10.1007/s00244-018-0580-4

D’Havé, H., Scheirs, J., Mubiana, V.K., Verhagen, R., Blust, R., De Coen, W., 2006. Non-destructive pollution exposure assessment in the European hedgehog (*Erinaceus europaeus*): II. Hair and spines as indicators of endogenous metal and As concentrations. Environmental Pollution 142, 438–448. 10.1016/j.envpol.2005.10.021

Douhard, F., Gaillard, J.-M., Pellerin, M., Jacob, L., Lemaître, J.-F., 2017. The cost of growing large: Costs of post-weaning growth on body mass senescence in a wild mammal. Oikos 126, 1329–1338. 10.1111/oik.04421

Draghi, S., Agradi, S., Riva, F., Tarhan, D., Bilgiç, B., Dokuzeylül, B., Ercan, A.M., Or, M.E., Brecchia, G., Vigo, D., Arioli, F., Di Cesare, F., Curone, G., 2023. Roe deer (*Capreolus capreolus*) hair as a bioindicator for the environmental presence of toxic and trace elements. Toxics 11, 49. 10.3390/toxics11010049

Fan, Y., Zhao, X., Yu, J., Xie, J., Li, C., Liu, D., Tang, C., Wang, C., 2020. Lead-induced oxidative damage in rats/mice: A meta-analysis. J Trace Elem Med Biol 58, 126443. 10.1016/j.jtemb.2019.126443

Finkel, T., Holbrook, N.J., 2000. Oxidants, oxidative stress and the biology of ageing. Nature 408, 239–247. 10.1038/35041687

Flerov, K.K., 1952. Fauna of USSR, Mammals Vol. 1, No. 2. Musk Deer and Deer. USSR Academia Science.

Gaillard, J.-M., Delorme, D., Jean-Marie, B., Van Laere, G., Boisaubert, B., Pradel, R., 1993. Roe Deer Survival Patterns: A Comparative Analysis of Contrasting Populations. Journal of Animal Ecology 62, 778–791. 10.2307/5396

Gaillard, J.-M., Mark Hewison, A.J., Klein, F., Plard, F., Douhard, M., Davison, R., Bonenfant, C., 2013. How does climate change influence demographic processes of widespread species? Lessons from the comparative analysis of contrasted populations of roe deer. Ecology Letters 16, 48–57. 10.1111/ele.12059

García, M.H. de M., Moreno, D.H., Rodríguez, F.S., Beceiro, A.L., Álvarez, L.E.F., López, M.P., 2011. Sex- and age-dependent accumulation of heavy metals (Cd, Pb and Zn) in liver, kidney and muscle of roe deer (*Capreolus capreolus*) from NW Spain. Journal of Environmental Science and Health, Part A 46, 109–116. 10.1080/10934529.2011.532422

Goutte, A., Barbraud, C., Meillère, A., Carravieri, A., Bustamante, P., Labadie, P., Budzinski, H., Delord, K., Cherel, Y., Weimerskirch, H., Chastel, O., 2014. Demographic consequences of heavy metals and persistent organic pollutants in a vulnerable long-lived bird, the wandering albatross. Proceedings of the Royal Society B: Biological Sciences 281, 20133313. 10.1098/rspb.2013.3313

Goutte, A., Meillère, A., Barbraud, C., Budzinski, H., Labadie, P., Peluhet, L., Weimerskirch, H., Delord, K., Chastel, O., 2018. Demographic, endocrine and behavioral responses to mirex in the South polar skua. Science of The Total Environment 631–632, 317–325. 10.1016/j.scitotenv.2018.02.326

Govind, P., 2014. Heavy metals causing toxicity in animals and fishes. Research Journal of Animal, Veterinary and Fishery Sciences 2, 17–23.

Halliwell, B., Gutteridge, J.M.C., 2015. Free radicals in biology and medicine. Oxford University Press.

Herrada, A., Bariod, L., Saïd, S., Rey, B., Bidault, H., Bollet, Y., Chabot, S., Débias, F., Duhayer, J., Pardonnet, S., Pellerin, M., Fanjul, J.-B., Rousset, C., Fritsch, C., Crini, N., Scheifler, R., Bourgoin, G., Vuarin, P., 2024a. Minor and trace element concentrations in roe deer hair: A non-invasive method to define reference values in wildlife. Ecological Indicators 159, 111720. 10.1016/j.ecolind.2024.111720

Herrada, A., Vuarin, P., Débias, F., Gache, A., Veber, P., Pellerin, M., Cheynel, L., Lemaître, J.-F., Gilot-Fromont, E., Rey, B., 2024b. Haemolysis overestimates plasma oxidative stress biomarkers in free-ranging roe deer. Comparative Biochemistry and Physiology Part A: Molecular & Integrative Physiology 298, 111750. 10.1016/j.cbpa.2024.111750

Iavicoli, I., Fontana, L., Bergamaschi, A., 2009. The effects of metals as endocrine disruptors. Journal of Toxicology and Environmental Health, Part B 12, 206–223. 10.1080/10937400902902062

Isaksson, C., 2020. Urban ecophysiology: Beyond costs, stress and biomarkers. J Exp Biol 223, jeb203794. 10.1242/jeb.203794

Isaksson, C., 2010. Pollution and its impact on wild animals: A meta-analysis on oxidative stress. EcoHealth 7, 342–350. 10.1007/s10393-010-0345-7

Johnson, E., Hornby, J., 1975. Seasonal changes of pelage in the roe deer *Capreolus capreolus* and its role in thermoregulation. Journal of Natural History 9, 619–628. 10.1080/00222937500770511

Kumar, A., Khushboo, Pandey, R., Sharma, B., 2020. Modulation of superoxide dismutase activity by mercury, lead, and arsenic. Biological Trace Element Research 196, 654–661. 10.1007/s12011-019-01957-3

Lazarus, M., Sergiel, A., Orct, T., Ferenčaković, M., Tariba Lovaković, B., Žunec, S., Rašić, D., Oster, E., Reljić, S., Macur, J., Huber, Đ., 2024. Apex carnivores coping with metal(loid) pollution and oxidative stress: Biological and environmental drivers of variation in kidney of European brown bear. Environmental Pollution 363, 125285. 10.1016/j.envpol.2024.125285

Linnell, J., Andersen, R., 1995. Site tenacity in roe deer: Short term effects of logging. Wildlife Society Bulletin 23, 31–36.

López-Otín, C., Blasco, M.A., Partridge, L., Serrano, M., Kroemer, G., 2023. Hallmarks of aging: An expanding universe. Cell 186, 243–278. 10.1016/j.cell.2022.11.001

Losdat, S., Helfenstein, F., Gaude, B., Richner, H., 2011. Reproductive effort transiently reduces antioxidant capacity in a wild bird. Behavioral Ecology 22, 1218–1226. 10.1093/beheco/arr116

Lüdecke, D., Ben-Shachar, M., Patil, I., Waggoner, P., Makowski, D., 2021. performance: An R package for assessment, comparison and testing of statistical models. Journal of Open Source Software 6, 3139. 10.21105/joss.03139

Ludolphy, C., Kierdorf, U., Kierdorf, H., 2021. Lead concentrations in antlers of European roe deer (*Capreolus capreolus*) from an agricultural area in Northern Germany over a 119-year period - a historical biomonitoring study. Environmental Science and Pollution Research 28, 56069–56078. 10.1007/s11356-021-14538-6

Lushchak, V.I., 2014. Classification of oxidative stress based on its intensity. Experimental and Clinical Sciences Journal 13, 922–937.

Lushchak, V.I., 2011. Environmentally induced oxidative stress in aquatic animals. Aquatic Toxicology 101, 13–30. 10.1016/j.aquatox.2010.10.006

Martin, C., Capilla-Lasheras, P., Monaghan, P., Burraco, P., 2024. The impact of chemical pollution across major life transitions: a meta-analysis on oxidative stress in amphibians. Proc. R. Soc. B. 291. 10.1098/rspb.2024.1536

McLean, C.M., Koller, C.E., Rodger, J.C., MacFarlane, G.R., 2009. Mammalian hair as an accumulative bioindicator of metal bioavailability in Australian terrestrial environments. The Science of the Total Environment 407, 3588–3596. 10.1016/j.scitotenv.2009.01.038

Metcalfe, N.B., Alonso-Alvarez, C., 2010. Oxidative stress as a life-history constraint: the role of reactive oxygen species in shaping phenotypes from conception to death. Functional Ecology 24, 984–996. 10.1111/j.1365-2435.2010.01750.x

Météo-France [WWW Document], 2023. URL https://www.data.gouv.fr/fr/organizations/meteo-france/ (accessed 2.11.25).

Montes, I., Newman, C., Mian, R., Macdonald, D.W., 2011. Radical health: Ecological corollaries of body condition, transport stress and season on plasma antioxidant capacity in the European badger. Journal of Zoology 284, 114–123. 10.1111/j.1469-7998.2010.00787.x

Moreau, J., Monceau, K., Crépin, M., Tochon, F.D., Mondet, C., Fraikin, M., Teixeira, M., Bretagnolle, V., 2021. Feeding partridges with organic or conventional grain triggers cascading effects in life-history traits. Environmental Pollution 278, 116851. 10.1016/j.envpol.2021.116851

Murphy, M.P., Bayir, H., Belousov, V., Chang, C.J., Davies, K.J.A., Davies, M.J., Dick, T.P., Finkel, T., Forman, H.J., Janssen-Heininger, Y., Gems, D., Kagan, V.E., Kalyanaraman, B., Larsson, N.-G., Milne, G.L., Nyström, T., Poulsen, H.E., Radi, R., Van Remmen, H., Schumacker, P.T., Thornalley, P.J., Toyokuni, S., Winterbourn, C.C., Yin, H., Halliwell, B., 2022. Guidelines for measuring reactive oxygen species and oxidative damage in cells and in vivo. Nature Metabolism 4, 651–662. 10.1038/s42255-022-00591-z

Nzengue, Y., Candéias, S.M., Sauvaigo, S., Douki, T., Favier, A., Rachidi, W., Guiraud, P., 2011. The toxicity redox mechanisms of cadmium alone or together with copper and zinc homeostasis alteration: Its redox biomarkers. Journal of Trace Elements in Medicine and Biology 25, 171–180. 10.1016/j.jtemb.2011.06.002

Ouyang, J.Q., Lendvai, Á.Z., Moore, I.T., Bonier, F., Haussmann, M.F., 2016. Do hormones, telomere lengths, and oxidative stress form an integrated phenotype? A case study in free-living tree swallows. Integrative and Comparative Biology 56, 138–145. 10.1093/icb/icw044

Pellerin, M., Calenge, C., Saïd, S., Gaillard, J.-M., Fritz, H., Duncan, P., Van Laere, G., 2010. Habitat use by female western roe deer (*Capreolus capreolus*): Influence of resource availability on habitat selection in two contrasting years. Canadian Journal of Zoology 88, 1052–1062. 10.1139/Z10-070

Pettorelli, N., Gaillard, J.-M., Mysterud, A., Duncan, P., Chr. Stenseth, N., Delorme, D., Van Laere, G., Toïgo, C., Klein, F., 2006. Using a proxy of plant productivity (NDVI) to find key periods for animal performance: The case of roe deer. Oikos 112, 565–572. 10.1111/j.0030-1299.2006.14447.x

Pokorny, B., Ribaric-Lasnik, C., 2002. Seasonal variability of mercury and heavy metals in roe deer (*Capreolus capreolus*) kidney. Environmental Pollution (Barking, Essex: 1987) 117, 35–46. 10.1016/s0269-7491(01)00161-0

R core Team, 2021. R: A language and environment for statistical computing. R Foundation for Statistical Computing, Vienna, Austria [WWW Document]. URL https://www.r-project.org/

Rashed, M.N., Soltan, M.E., 2005. Animal hair as biological indicator for heavy metal pollution in urban and rural areas. Environmental monitoring and assessment 110, 41–53. 10.1007/s10661-005-6288-8

Rattner, B.A., 2009. History of wildlife toxicology. Ecotoxicology 18, 773–783. 10.1007/s10646-009-0354-x

Rodríguez-Estival, J., de la Lastra, J.M.P., Ortiz-Santaliestra, M.E., Vidal, D., Mateo, R., 2013. Expression of immunoregulatory genes and its relationship to lead exposure and lead-mediated oxidative stress in wild ungulates from an abandoned mining area. Environmental Toxicology and Chemistry 32, 876–883. 10.1002/etc.2134

Rodríguez-Estival, J., García-de Blas, E., Smits, J.E.G., 2016. Oxidative stress biomarkers indicate sublethal health effects in a sentinel small mammal species, the deer mouse (*Peromyscus maniculatus*), on reclaimed oil sands areas. Ecological Indicators 62, 66–75. 10.1016/j.ecolind.2015.11.019

Rodríguez-Estival, J., Martinez-Haro, M., Monsalve-González, L., Mateo, R., 2011. Interactions between endogenous and dietary antioxidants against Pb-induced oxidative stress in wild ungulates from a Pb polluted mining area. Science of The Total Environment 409, 2725–2733. 10.1016/j.scitotenv.2011.04.010

Saaristo, M., Brodin, T., Balshine, S., Bertram, M.G., Brooks, B.W., Ehlman, S.M., McCallum, E.S., Sih, A., Sundin, J., Wong, B.B.M., Arnold, K.E., 2018. Direct and indirect effects of chemical contaminants on the behaviour, ecology and evolution of wildlife. Proceedings of the Royal Society B: Biological Sciences 285, 20181297. 10.1098/rspb.2018.1297

Saïd, S., Gaillard, J.-M., Widmer, O., Débias, F., Bourgoin, G., Delorme, D., Roux, C., 2009. What shapes intra-specific variation in home range size? A case study of female roe deer. Oikos 118, 1299–1306. 10.1111/j.1600-0706.2009.17346.x

Sempéré, A.J., Sokolov, V.E., Danilkin, A.A., 1996. Capreolus capreolus. Mammalian Species 1–9.

Shahid, M., Natasha, Dumat, C., Niazi, N.K., Xiong, T.T., Farooq, A.B.U., Khalid, S., 2019. Ecotoxicology of heavy metal(loid)-enriched particulate matter: Foliar accumulation by plants and health impacts, in: de Voogt, P. (Ed.), Reviews of Environmental Contamination and Toxicology, Reviews of Environmental Contamination and Toxicology. Springer International Publishing, pp. 65–113. 10.1007/398_2019_38

Shore, R.F., Rattner, B.A., 2001. Ecotoxicology of wild mammals. Wiley.

Soetan, K.O., Olaiya, C.O., Oyewole, O.E., 2010. The importance of mineral elements for humans, domestic animals and plants: A review. African Journal of Food Science 4, 200–222.

Sokolova, I., Lannig, G., 2008. Interactive effects of metal pollution and temperature on metabolism in aquatic ectotherms: Implications of global climate change. Climate Research 37, 181–201. 10.3354/cr00764

Strandgaard, H., 1972. The roe deer (*Capreolus capreolus*) population at Kalø and the factors regulating its size. Danish Review of Game Biology 7, 205.

Tapiero, H., Townsend, D.M., Tew, K.D., 2003. The antioxidant role of selenium and seleno-compounds. Biomedicine & pharmacotherapy 57, 134. 10.1016/s0753-3322(03)00035-0

Tchounwou, P.B., Yedjou, C.G., Patlolla, A.K., Sutton, D.J., 2012. Heavy metal toxicity and the environment, in: Luch, A. (Ed.), Molecular, Clinical and Environmental Toxicology, Experientia Supplementum. Springer Basel, Basel, pp. 133–164. 10.1007/978-3-7643-8340-4_6

Tête, N., Afonso, E., Crini, N., Drouhot, S., Prudent, A.-S., Scheifler, R., 2014. Hair as a noninvasive tool for risk assessment: Do the concentrations of cadmium and lead in the hair of wood mice (*Apodemus sylvaticus*) reflect internal concentrations? Ecotoxicology and Environmental Safety 108, 233–241. 10.1016/j.ecoenv.2014.07.010

Thioulouse, J., Dray, S., Dufour, A.-B., Siberchicot, A., Jombart, T., Pavoine, S., 2018. Multivariate analysis of ecological data with ade4, Springer New York, NY. ed.

USEPA, 2000. Guidance for data quality assessment.

Valko, M., Morris, H., Cronin, M., 2005. Metals, toxicity and oxidative stress. Current Medicinal Chemistry 12, 1161–1208. 10.2174/0929867053764635

Vuarin, P., Rey, B., Herrada, A., Débias, F., Crini, N., Scheifler, R., Fritsch, C., Lemaître, J.-F., Saïd, S., 2025. Exposure to toxic trace metals is negatively associated with testis mass in wild male roe deer. Environmental Pollution 126825. 10.1016/j.envpol.2025.126825

Wirth, J.J., Mijal, R.S., 2010. Adverse effects of low level heavy metal exposure on male reproductive function. Systems Biology in Reproductive Medicine 56, 147–167. 10.3109/19396360903582216

Xiao, W., Zhang, Y., Chen, X., Sha, A., Xiong, Z., Luo, Y., Peng, L., Zou, L., Zhao, C., Li, Q., 2024. The easily overlooked effect of global warming: Diffusion of heavy metals. Toxics 12, 400. 10.3390/toxics12060400

Xu, M., Rui, D., Yan, Y., Xu, S., Niu, Q., Feng, G., Wang, Y., Li, S., Jing, M., 2017. Oxidative damage induced by arsenic in mice or rats: A systematic review and meta-analysis. Biol Trace Elem Res 176, 154–175. 10.1007/s12011-016-0810-4

Zoroddu, M.A., Aaseth, J., Crisponi, G., Medici, S., Peana, M., Nurchi, V.M., 2019. The essential metals for humans: A brief overview. Journal of Inorganic Biochemistry 195, 120–129. 10.1016/j.jinorgbio.2019.03.013

