## Supplementary material for "Unravelling the association between non-essential elements and redox homeostasis in wild roe deer": Table S1

1. **Limits of quantification and treatment of values below the LOQ**

| **Table S1.** Limits of quantification (LOQ) and proportion of element concentrations below the LOQ in roe deer hair. For each element, the median LOQ and its range across samples are reported. Values below the LOQ were assigned a concentration equal to half of the corresponding LOQ (LOQ/2) for statistical analyses. | | | | |
| --- | --- | --- | --- | --- |
|  |  | **Unit** | **LOQ** | **% values below LOQ** |
|  |  |  | **median (min–max)** |  |
| Non-essential elements | Al | μg/g | 1.54 (1.26–1.97) | 0 |
|  | As | ng/g | 7.19 (5.75–11.9) | 11.2 |
|  | Cd | ng/g | 4.99 (2.22–19.4) | 25.7 |
|  | Hg | ng/g | 7.96 (3.66–9.90) | 10.3 |
|  | Pb | μg/g | 0.00537 (0.00355–0.00556) | 0 |
|  | Sb | ng/g | 2.54 (1.61–5.69) | 15.4 |
|  | Sr | μg/g | 0.136 (0.0588–0.222) | 18.0 |
|  | Ti | μg/g | 0.0175 (0.00405–0.0186) | 0 |
|  | Tl | ng/g | 0.212 (0.0934–0.301) | 12.0 |
| Essential elements | Ca | μg/g | 14.7 (13.2–17.7) | 0 |
|  | Co | ng/g | 2.46 (1.61–3.16) | 0 |
|  | Cr | ng/g | 13.0 (7.49–24.7) | 1.3 |
|  | Cu | μg/g | 0.0377 (0.0244–0.0552) | 0 |
|  | Fe | μg/g | 0.140 (0.0189–0.952) | 0 |
|  | K | μg/g | 65.1 (57.5–118) | 0 |
|  | Mg | μg/g | 47.6 (38.2–62.2) | 0 |
|  | Mn | μg/g | 0.0136 (0.00850–0.0336) | 0 |
|  | Mo | ng/g | 4.74 (1.18–7.86) | 0.6 |
|  | Na | μg/g | 18.2 (13.6–19.1) | 0 |
|  | Ni | ng/g | 31.6 (14.5–35.8) | 1.3 |
|  | Se | μg/g | 0.0177 (0.00425–0.0918) | 0.3 |
|  | Zn | μg/g | 0.324 (0.0771–1.40) | 0 |

2. **Raw data on oxidative stress biomarkers**

Raw values of oxidative stress biomarkers measured in roe deer plasma (dROMs, MDA, Carbonyls, OXY, SOD) or erythrocytes (GSH, GSH:GSSG) are presented in Table S1. For each biomarker, the mean value, the median one and the range of concentrations (i.e. min and max values) before scaling are provided.

| **Table S2**. Raw values of oxidative stress biomarkers measured in roe deer plasma or erythrocytes from samples collected between 2016 and 2019. dROMS: reactive oxygen metabolites, MDA: malondialdehyde, Carbonyls: protein carbonyl content, OXY: total antioxidant capacity of plasma, SOD: superoxide dismutase, GSH: reduced glutathione, GSSG: oxidised glutathione. | | | | | |
| --- | --- | --- | --- | --- | --- |
| **Type of marker** | **Marker** | **Unit** | **Mean ± sd** | **Median** | **Range (min-max)** |
| Oxidative  damage | **dROMs** (*n* = 636) | mg/dL | 8.18 ± 2.59 | 8.16 | 0.00 - 20.0 |
|  | **MDA** (*n* = 632) | µM | 12.1 ± 6.60 | 10.0 | 2.59 - 42.7 |
|  | **Carbonyls** (*n* = 627) | nmol/mg | 0.97 ± 0.29 | 0.93 | 0.40 - 3.68 |
| Antioxidant  defences | **OXY** (*n* = 636) | µmol/mL | 188 ± 39.9 | 182 | 54.3 - 360 |
|  | **SOD** (*n* = 632) | % inhibition rate | 70.6 ± 8.68 | 70.2 | 44.8 - 95.4 |
|  | **GSH** (*n* = 245) | µmol/L | 18.3 ± 7.57 | 17.3 | 3.50 - 49.2 |
| Oxidative status | **GSH:GSSG** (*n* = 245) | AU | 9.57 ± 8.21 | 6.74 | 1.32 - 66.3 |

3. **Effects of weather conditions on hair non-essential element concentrations**

Weather conditions may affect non-essential elements uptake by plants and animals, and consequently accumulation patterns. Indeed, element bioavailability is known to be influenced by the chemical and mineralogical characteristics of the soil, that are tightly linked to the weather. More specifically, it has been found that the bioaccumulation of non-essential elements in plants and animals is stronger in warm and dry environments (Xiao et al., 2024). Interestingly, Pokorny and Ribaric-Lasnik (Pokorny and Ribaric-Lasnik, 2002) highlighted a strong seasonal difference in Hg and Pb concentrations in roe deer kidneys from Slovenia, with significantly higher levels in late summer and early autumn than in the other two tested seasons. These results were partly attributed to increased food intake during this period of the year, as well as a dietary shift towards element-accumulating mushrooms.

As roe deer captures happen in winter, when the winter coat is fully set since roe deer undergo an autumnal moult, we hypothesised that the storage of non-essential elements happened from September to December, and was therefore completed by the time of sampling. Hence, we retrieved daily temperatures and rainfall from the nearest locations to Chizé and Trois-Fontaines forests using Météo-France data (Direction interministérielle du numérique (DINUM), 2023). We averaged temperatures, as well as summed up cumulative rainfall, over this 3-month period for each capture year and each population, representing a total of 8 values. We then computed linear models where either PCT1 or PCT2 coordinates were set as the dependent variable, and mean ambient temperatures, cumulative rainfall and population were set as explanatory variables. Population was included to account for fixed differences between the two study sites.

Results (Fig. S2) highlighted weak but significant positive correlations between cumulative rainfall and PCT1 and PCT2 coordinates (β = 0.008 [0.005 ; 0.011], p < 0.001 and β = 0.008 [0.007 ; 0.009], p < 0.001, respectively), but no significant correlation between mean temperature and PCT1 or PCT2 coordinates since the temperature association was not distinguishable from site-level differences. Hence, higher cumulative rainfall seems to accentuate non-essential element accumulation in roe deer hair. Note that, however, weather conditions have only small effects on element accumulation during hair growth, and that the exact mechanisms underlying the correlations remain unknown.

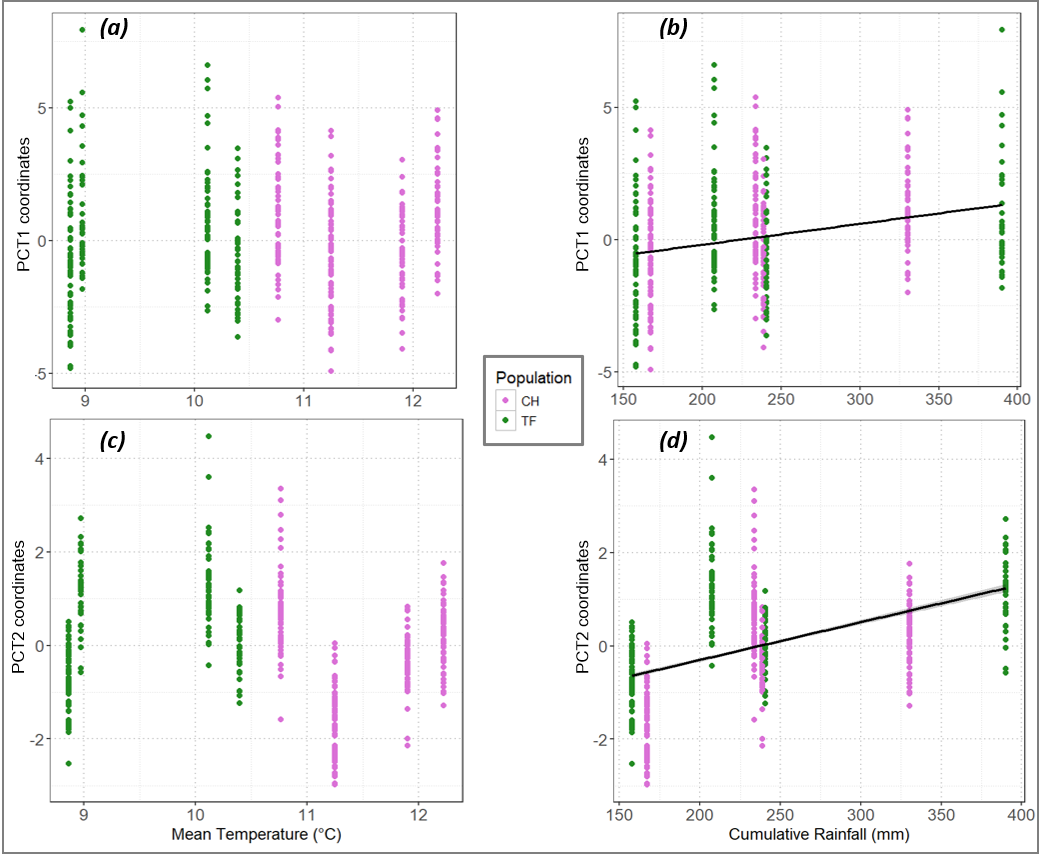

**Figure S1.** Effects of mean ambient temperature and cumulative rainfall on PCT1 and PCT2 coordinates. Points represent raw data, with lilac dots for individuals from Chizé (CH) and green dots for individuals from Trois-Fontaines (TF). Black lines with shaded areas represent the predictions from linear models, and are only shown when the relationships are significant. Relationships between mean ambient temperature and (a) PCT1 or (c) PCT2 coordinates, and between cumulative rainfall and (b) PCT1 or (d) PCT2 coordinates.

**Direction interministérielle du numérique (DINUM)** (2023). Météo-France.

**Pokorny, B. and Ribaric-Lasnik, C.** (2002). Seasonal variability of mercury and heavy metals in roe deer (*Capreolus capreolus*) kidney. *Environmental Pollution (Barking, Essex: 1987)* **117**, 35–46.

**Xiao, W., Zhang, Y., Chen, X., Sha, A., Xiong, Z., Luo, Y., Peng, L., Zou, L., Zhao, C. and Li, Q.** (2024). The easily overlooked effect of global warming: Diffusion of heavy metals. *Toxics* **12**, 400.

4. **Additional Principal Component Analyses outputs**

Since our dataset includes repeated measurements on the same individuals, and individuals from two contrasted populations, we also performed PCAs on non-essential (Fig. S2) and essential (Fig. S3) elements excluding those repeated measurements and analysing both populations separately. For the analysis based on single measurements, one sample was randomly selected for each individual with multiple records.

These additional analyses revealed that results remain consistent regardless of the inclusion of repeated measurements (Fig. S2a; Fig. S3a) or whether populations were analysed together or separately (Fig. S2b,c; Fig. S3b,c).

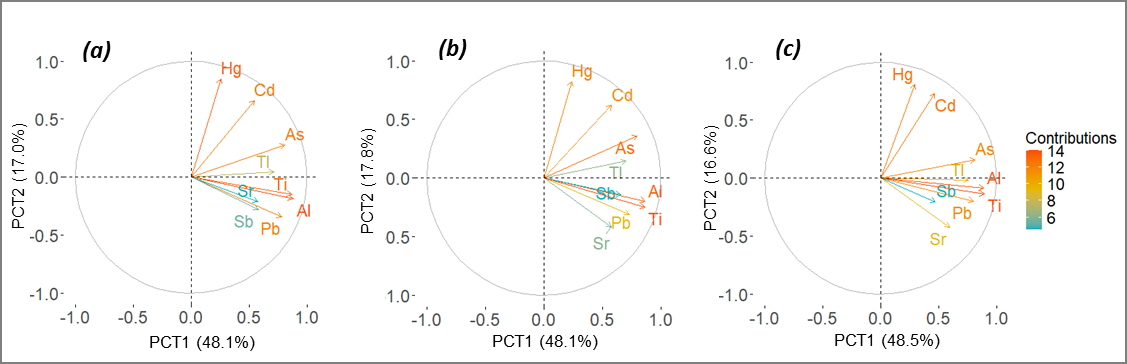

**Figure S2.** First factorial plan of the Principal Component Analysis (PCA) based on 9 non-essential elements, showing variable contributions for (a) single measurements (n = 434), (b) Chizé (n = 362), and for (c) Trois-Fontaines (n = 287) populations. Colours represent the contribution of the variables on the first axis.

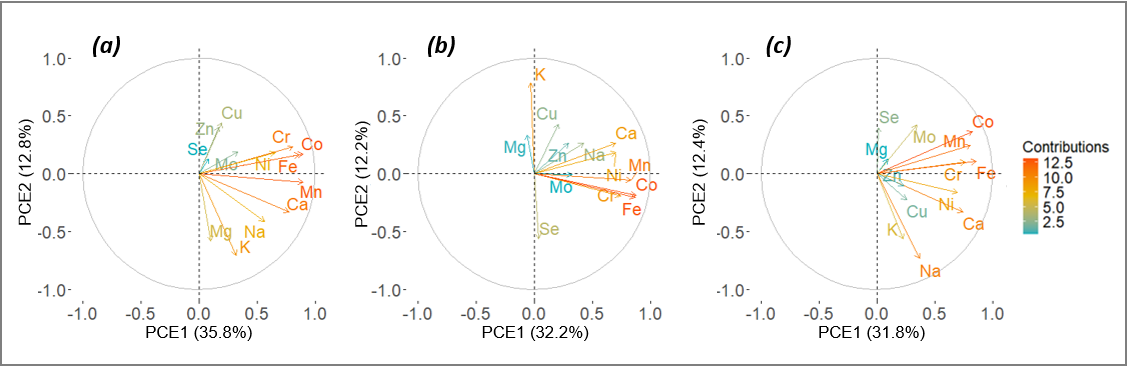

**Figure S3.** First factorial plan of the Principal Component Analysis (PCA) based on 13 essential elements, showing variable contributions for (a) single measurements (n = 434), (b) Chizé (n = 362), and for (c) Trois-Fontaines (n = 287) populations. Colours represent the contribution of the variables on the first axis. CHANGER

5. **Retained models within 2 ΔAICc and outputs of the best-supported models**

The set of models retained within 2 ΔAICc of the best-supported model, as well as the corresponding model outputs, are presented in Table S3 and S4, respectively. The most parsimonious model was then selected for Table S4, and for graphical presentation (see Figs. 3 & 4). When models are equivalent (same number of parameters and within 2 ΔAICc of the best-supported model), these models may be considered in the interpretation of results.

| **Table S3.** Retained models within 2 ΔAICc describing markers of oxidative damage (dROMs, MDA, Carbonyls), antioxidant defences (OXY, SOD, GSH) and oxidative status (GSH:GSSG) as a function of indices of non-essential element exposure (PCT 1 or 2), indices of essential element exposure (PCE 1 or 2), their interaction (PCT:PCE), cumulative rainfall (Rain) and its interaction with PCT (PCT:Rain), mean ambient temperature (Temp) and its interaction with PCT (PCT:Temp), age, body mass, plasma haemoglobin levels (Haemoglobin), population (Pop), sex and year. For sex and population variables, intercepts include females and individuals from Chizé, respectively. Models are sorted by increasing ΔAICc. The best-supported models are highlighted in bold. Full models are added for comparison. dROMS: reactive oxygen metabolites, MDA: malondialdehyde, Carbonyls: protein carbonyl content, OXY: total antioxidant capacity of plasma, SOD: superoxide dismutase, GSH: reduced glutathione, GSSG: oxidised glutathione. | | | | | | |
| --- | --- | --- | --- | --- | --- | --- |
| **Biomarker** | **Model number** | **Model structure** | **Parameters** | **AICc** | **ΔAICc** | **AICc weight** |
| dROMs | 535696 | PCT2*Rain + PCT2*Temp + Age + Body mass + Haemoglobin + Year | 12 | 1760.5 | 0.00 | 0.334 |
|  | 527504 | PCT2*Rain + Age + Body mass + Haemoglobin + Year | 11 | 1760.7 | 0.15 | 0.310 |
|  | 533648 | PCT2*Rain + PCT2*Temp + Age + Body mass + Haemoglobin | 11 | 1761.7 | 1.12 | 0.191 |
|  | **525456** | **PCT2*Rain + Temp + Age + Body mass + Haemoglobin** | **10** | **1761.9** | **1.41** | **0.165** |
|  | 1048576 | *Full model* | 23 | 1773.8 | 13.24 |  |
| MDA | **10635** | **PCT2*Temp + Haemoglobin + Pop + Year** | **8** | **923.3** | **0.00** | **1.000** |
|  | 1048576 | *Full model* | 22 | 944.4 | 21.1 |  |
| Carbonyls | **172224** | **PCT2*PCE1 + PCT2*PCE2 + PCT2*Temp + Age + Body mass + Haemoglobin** | **12** | **1131.0** | **0.00** | **0.400** |
|  | 172476 | PCT2*PCE1 + PCT2*PCE2 + PCT2*Temp + Age + Haemoglobin + Population | 12 | 1131.3 | 0.36 | 0.334 |
|  | 1048576 | *Full model* | 22 | 1131.8 | 0.81 | 0.266 |
| OXY | 789959 | PCT1*Rain + PCT2*Rain + Body mass + Haemoglobin + Pop + Year | 12 | 1636.2 | 0.00 | 0.279 |
|  | 789957 | PCT1*Rain + PCT2*Rain + Body mass + Pop + Year | 11 | 1636.4 | 0.14 | 0.259 |
|  | 527815 | PCT1 + PCT2*Rain + Body mass + Haemoglobin + Pop + Year | 11 | 1636.9 | 0.65 | 0.201 |
|  | **527813** | **PCT1 + PCT2*Rain + Body mass + Pop + Year** | **10** | **1637.4** | **1.14** | **0.157** |
|  | 527767 | PCT2*Rain + PCE1 + Body mass + Haemoglobin + Pop + Year | 11 | 1638.2 | 1.98 | 0.104 |
|  | 1048576 | *Full model* | 23 | 1652.3 | 16.08 |  |
| SOD | **528031** | **PCT2*Rain + PCE1 + Temp + Body mass + Haemoglobin + Sex + Year** | **11** | **968.8** | **0.00** | **0.359** |
|  | 528335 | PCT1 + PCT2*Rain + Temp + Body mass + Haemoglobin + Population + Sex + Year | 12 | 968.9 | 0.08 | 0.346 |
|  | 528079 | PCT1 + PCT2*Rain + Temp + Body mass + Haemoglobin + Sex + Year | 11 | 969.2 | 0.40 | 0.295 |
|  | 1048576 | *Full model* | 22 | 985.2 | 16.4 |  |
| GSH | **2053** | **Body mass + Year** | **4** | **564.9** | **0.00** | **1.000** |
|  | 1048576 | *Full model* | 22 | 594.1 | 29.14 |  |
| GSH:GSSG | 953597 | PCT1*Rain + PCT2*PCE1 + PCT2*PCE2 + PCT2*Rain + Temp + Body mass + Year | 14 | 486.3 | 0.00 | 0.327 |
|  | 429309 | PCT1*Rain + PCT2*PCE1 + PCT2*PCE2 + Temp + Body mass + Year | 13 | 486.7 | 0.42 | 0.264 |
|  | 170237 | PCT1*Temp + PCT2*PCE1 + PCT2*PCE2 + Body mass + Year | 12 | 487.2 | 0.91 | 0.207 |
|  | **166141** | **PCT1 + PCT2*PCE1 + PCT2*PCE2 + Temp + Body mass + Year** | **11** | **487.2** | **0.96** | **0.203** |
|  | 1048576 | *Full model* | 22 | 499.9 | 13.63 |  |

| **Table S4**. Parameter estimates of linear (mixed-effects) models describing markers of oxidative damage (dROMs, MDA, Carbonyls), antioxidant defences (OXY, SOD, GSH) and oxidative status (GSH:GSSG) as a function of indices of non-essential element exposure (PCT 1 or 2), indices of essential element exposure (PCE 1 or 2), their interaction (PCT:PCE), cumulative rainfall (Rain) and its interaction with PCT (PCT:Rain), mean ambient temperature (Temp) and its interaction with PCT (PCT:Temp), age, body mass, plasma haemoglobin levels (Haemoglobin), population (Pop), sex and year. For sex and population variables, intercepts include females and individuals from Chizé, respectively. Parameter estimates are given with 95% confidence intervals (95% CI). Estimates whose 95% CI do not cross zero are displayed in bold. dROMS: reactive oxygen metabolites, MDA: malondialdehyde, Carbonyls: protein carbonyl content, OXY: total antioxidant capacity of plasma, SOD: superoxide dismutase, GSH: reduced glutathione, GSSG: oxidised glutathione, SD: standard deviation. | | | | | | | | | | | | | | |
| --- | --- | --- | --- | --- | --- | --- | --- | --- | --- | --- | --- | --- | --- | --- |
| **Fluid analysed** | **Plasma** | | | | | | | | | | **Erythrocytes** | | | |
| **Type of marker** | **Oxidative damage** | | | | | | **Antioxidant defences** | | | | | | **Oxidative status** | |
| **Variables** | **dROMs** (*n* = 636) | | **MDA** (*n* = 426) | | **Carbonyls** (*n* = 424) | | **OXY** (*n* = 636) | | **SOD** (*n* = 427) | | **GSH** (*n* = 218) | | **GSH:GSSG** (*n* = 218) | |
| **Fixed effects** | Estimate | 95% CI | Estimate | 95% CI | Estimate | 95% CI | Estimate | 95% CI | Estimate | 95% CI | Estimate | 95% CI | Estimate | 95% CI |
| Intercept | -2.21e^-03^ | [-0.08 ; 0.08] | **-0.19** | **[-0.29 ; -0.09]** | 0.01 | [-0.08 ; 0.10] | **0.16** | **[0.06 ; 0.26]** | **-0.13** | **[-0.23 ; -0.04]** | **0.40** | **[0.24 ; 0.56]** | **0.41** | **[0.23 ; 0.59]** |
| PCT1 |  |  |  |  |  |  | **0.05** | **[0.02 ; 0.09]** |  |  |  |  | **0.22** | **[0.10 ; 0.33]** |
| PCT2 | **0.07** | **[0.01 ; 0.13]** | **0.13** | **[0.07 ; 0.20]** | **0.17** | **[0.10 ; 0.25]** | **0.13** | **[0.06 ; 0.20]** | **-0.26** | **[-0.33 ; -0.19]** |  |  | -0.01 | [-0.12 ; 0.10] |
| PCE1 |  |  |  |  | **0.06** | **[0.02 ; 0.10]** |  |  | **-0.05** | **[-0.09 ; -0.01]** |  |  | **-0.17** | **[-0.27 ; -0.07]** |
| PCE2 |  |  |  |  | **0.11** | **[0.03 ; 0.18]** |  |  |  |  |  |  | **-0.12** | **[-0.22 ; -0.03]** |
| PCT1:PCE1 |  |  |  |  |  |  |  |  |  |  |  |  |  |  |
| PCT1:PCE2 |  |  |  |  |  |  |  |  |  |  |  |  |  |  |
| PCT2:PCE1 |  |  |  |  | **0.04** | **[0.01 ; 0.08]** |  |  |  |  |  |  | **0.06** | **[0.02 ; 0.10]** |
| PCT2:PCE2 |  |  |  |  | **0.08** | **[0.02 ; 0.15]** |  |  |  |  |  |  | **0.09** | **[0.03 ; 0.16]** |
| Rain | 0.04 | [-0.05 ; 0.12] |  |  |  |  | 0.06 | [-0.02 ; 0.14] | 2.42e^-03^ | [-0.09 ; 0.08] |  |  |  |  |
| PCT1:Rain |  |  |  |  |  |  |  |  |  |  |  |  |  |  |
| PCT2:Rain | **0.06** | **[0.01 ; 0.12]** |  |  |  |  | **0.07** | **[0.02 ; 0.12]** | **-0.07** | **[-0.13 ; -0.02]** |  |  |  |  |
| Temp | -0.07 | [-0.15 ; 0.01] | **0.23** | **[0.14 ; 0.31]** | -0.02 | [-0.12 ; 0.07] |  |  | **-0.12** | **[-0.21 ; -0.04]** |  |  | **0.15** | **[0.04 ; 0.26]** |
| PCT1:Temp |  |  |  |  |  |  |  |  |  |  |  |  |  |  |
| PCT2:Temp |  |  | **0.12** | **[0.06 ; 0.18]** | **0.17** | **[0.09 ; 0.25]** |  |  |  |  |  |  |  |  |
| Age | **-0.23** | **[-0.33 ; -0.13]** |  |  | **-0.17** | **[-0.29 ; -0.06]** |  |  |  |  |  |  |  |  |
| Body mass | **0.26** | **[0.16 ; 0.36]** |  |  | **0.13** | **[0.01 ; 0.25]** | **0.16** | **[0.08 ; 0.23]** | **-0.15** | **[-0.22 ; -0.07]** | **0.16** | **[0.05 ; 0.28]** | **-0.17** | **[-0.27 ; -0.07]** |
| Haemoglobin | **0.09** | **[0.02 ; 0.17]** | **0.17** | **[0.10 ; 0.25]** | **0.29** | **[0.19 ; 0.38]** |  |  | **0.47** | **[0.39 ; 0.54]** |  |  |  |  |
| Pop (TF) |  |  | **0.39** | **[0.23 ; 0.56]** |  |  | **-0.38** | **[-0.54 ; -0.22]** |  |  |  |  |  |  |
| Sex (M) |  |  |  |  |  |  |  |  | **0.31** | **[0.17 ; 0.45]** |  |  |  |  |
| Year |  |  | **0.52** | **[0.44 ; 0.60]** |  |  | **0.36** | **[0.27 ; 0.45]** | **-0.10** | **[-0.20 ; -0.01]** | **-0.48** | **[-0.62 ; -0.34]** | **-0.71** | **[-0.91 ; -0.52]** |
| **Random effects** | SD | |  |  |  |  | SD | |  |  |  |  |  |  |
| Individual ID | 0.43 | |  |  |  |  | 0.28 | |  |  |  |  |  |  |
| Marginal R² | 0.08 | | 0.49 | | 0.21 | | 0.25 | | 0.42 | | 0.20 | | 0.49 | |
| Conditional R² | 0.27 | |  |  |  |  | 0.32 | |  |  |  |  |  |  |
